# Chemo-omic pipeline enables discovery of prion synaptotoxic pathways and inhibitory drugs

**DOI:** 10.64898/2026.01.28.702331

**Authors:** Nhat T.T. Le, Robert C.C. Mercer, Cheng Fang, Aravind Sundaravadivelu, Adam T. Labadorf, Weiwei Lin, Julian Kwan, Benjamin Blum, Andrew Emili, David A. Harris

**Affiliations:** Department of Biochemistry & Cell Biology, Boston University Chobanian & Avedisian School of Medicine, Boston, MA 02118, USA; Bioinformatics Program, Boston University, Boston, MA 02215, USA; Department of Pharmacology, Physiology & Biophysics, Boston University Chobanian & Avedisian School of Medicine, Boston, MA 02118, USA; Department of Cancer Biology and Genetics, College of Medicine, The Ohio State University Columbus, Ohio 43210, USA; Department of Biochemistry & Molecular Genetics, College of Graduate Studies, Midwestern University, Glendale, Arizona 85308, USA; Annovis Bio Inc., Malvern, PA 19355, USA; Center for RNA Therapeutics, Houston Methodist Research Institute, Houston, Texas 77030, USA; Department of Biomedical Engineering, Division of Oncological Sciences, Knight Cancer Institute, Oregon Health and Science University, Portland, OR 97201, USA

## Abstract

Prion propagation, in which the cellular prion protein (PrP^C^) is conformationally converted into an infectious structure (PrP^Sc^), is now well understood. However, the molecular mechanism responsible for the neurotoxicity of prions remains unclear. Synaptic loss is one of the earliest events in both *in vivo* and *in vitro* models of prion disease. We previously developed a neuronal cell culture model to analyze the mechanisms of prion-induced synaptic degeneration in a physiologically relevant setting. Using this system, we showed that exposure of hippocampal neurons to PrP^Sc^ engages a NMDAR/p38 mitogen-activated protein kinase (MAPK) signaling pathway that results in rapid, PrP^C^-dependent loss of synaptic transmission and retraction of dendritic spines. To comprehensively identify the components of this synaptotoxic signaling pathway, we measured changes in the phosphoproteome and transcriptome of hippocampal neurons exposed to PrP^Sc^ while they were undergoing the process of dendritic spine retraction. We then used these data as input into the L1000 and P100 databases of transcriptomic and proteomic drug signatures, leading to the discovery of 17 compounds that were able to prevent PrP^Sc^-induced spine retraction. These compounds converged on three protein kinase targets: Ca^2+^/calmodulin-dependent protein kinase II (CaMKII), protein kinase C (PKC), and glycogen synthase kinase 3β (GSK3β). Using immunocytochemical staining, we confirmed that PrP^Sc^ treatment of hippocampal neurons induced phosphorylation of the three kinases and caused their rapid translocation to dendritic spines. Along with N-methyl-D-aspartate receptors (NMDARs) on the neuronal surface, which trigger an initial influx of Ca^2+^ in response to PrP^Sc^, these kinases constitute key nodes in a signaling network that mediates prion synaptotoxicity. Taken together, our results provide new insights into the mechanisms of prion neurotoxicity, and they identify novel molecular targets and inhibitory compounds that can be utilized for therapy of prion diseases.

**AUTHOR SUMMARY:** The mechanism by which prions propagate is now well established, but how they cause neurodegenerative changes is still uncertain. The earliest effects of prion infection occur at the level of the synapse, and we previously established an experimental system using cultured hippocampal neurons to assay prion synaptotoxicity. To search comprehensively for components of the synaptotoxic signaling pathway, we employed a novel, small-molecule discovery pipeline based on the transcriptomic and phosphoproteomic profiles of prion-treated neurons. This approach converged on inhibitors of three different protein kinases (Ca^2+^/calmodulin-dependent protein kinase II, protein kinase C, and glycogen synthase kinase 3β), which, along with N-methyl-D-aspartate receptors, constitute key nodes in a prion synaptotoxic signaling network that can be targeted for therapeutic benefit.

## INTRODUCTION

Prion diseases, or transmissible spongiform encephalopathies, are fatal neurodegenerative disorders of humans and animals. The key molecular event underlying these disorders is the conformational change of PrP^C^, an endogenous neuronal membrane glycoprotein, into a β-sheet rich conformation denoted PrP^Sc^ [1–3]. PrP knockout mice are completely resistant to prion infection [4, 5], suggesting that the disease phenotype is due primarily to a gain-of-function attributable to PrP^Sc^ or a related toxic species, rather than to a loss of a normal function of PrP^C^. While the mechanism of prion propagation is now relatively well understood, how PrP^Sc^ damages neurons has remained mysterious [6].

Synapse loss is an early feature shared by many neurodegenerative disorders [7, 8]. In prion diseases, synaptic loss begins well before symptoms are evident, and is likely to play a major role in the evolution of the disease [9]. However, little research has been done to determine the mechanism by which prions damage synapses, and to define the cellular pathways involved. Addressing these questions has been limited by the availability of *in vitro* model systems that can be interrogated at the cellular and molecular levels. Although several cell lines can be infected with prions, none display major prion-associated cytopathology [10]. Despite the importance of synaptic neurotoxicity in prion diseases, there is very little published literature on prion infection of cultured primary neurons [11–13].

To assay prion synaptotoxicity, we have developed a system in which neonatal mouse hippocampal neurons are cultured at low density on coverslips suspended over a feeder layer of astrocytes [14, 15]. This system allows precise morphological and electrophysiological characterization of synaptic contacts, as well as facile preparation of astrocyte-free samples for biochemical experiments. The hippocampal neurons display a rapid synaptotoxic response to prions, in which neurons retract their dendritic spines and exhibit alterations in synaptic transmission within 24 hours of exposure to PrP^Sc^. This system allows visualization of the earliest events in synaptic degeneration, before additional pathological changes and neuronal death ensue. Importantly, the changes in dendritic spines and synaptic transmission are entirely dependent on neuronal expression of PrP^C^ and occur without any observable reduction in neuronal viability.

Using this assay system, we have uncovered a prion synaptotoxic signaling pathway [15] that is initiated by rapid, PrP^Sc^-stimulated influx of Ca^2+^ ions via ionotropic NMDARs, followed by phosphorylation and activation of two downstream protein kinases: p38 mitogen-activated protein kinase (MAPK), which, in turn, activates MAPK-activated protein kinase (MAPKAP or MK). Stimulation of these kinases then leads to collapse of the F-actin cytoskeleton in dendritic spines, followed by spine retraction and loss of synaptic function. We worked out this pathway experimentally by using pharmacological and genetic methods to inhibit specific pathway components that we hypothesized might play a role.

In the present work, we sought an unbiased method to expand our map of the synaptotoxic signaling network and discover compounds to inhibit key nodes of the network. To accomplish this, we have applied a novel chemo-omic pipeline in which the transcriptomic and phosphoproteomic signatures of cultured neurons exposed to PrP^Sc^ were used to interrogate a set of public databases (L1000/P100) containing the corresponding signatures of a standard set of cell lines exposed to a large number of compounds [16–19]. This allowed us to identify compounds whose signatures were the inverse of those induced by PrP^Sc^. We confirmed that many of these compounds were potent inhibitors of PrP^Sc^-induced synaptic degeneration when tested in our hippocampal neuron culture system. These inhibitors converged on three key protein kinases (CaMKII, PKC, and GSK3β) that underwent phosphorylation and translocation to dendritic spines in response to PrP^Sc^. Together, our results define a kinase-dependent signaling network that mediates prion synaptotoxicity, and that provides a list of compounds and molecular targets for therapeutic intervention.

## RESULTS

### Time course of PrP^Sc^-induced synaptotoxicity

Previously, we showed that treatment of cultured hippocampal neurons for 24 hours with purified PrP^Sc^, but not control preparations, induced retraction of dendritic spines and impairment of synaptic transmission, effects that were entirely PrP^C^-dependent [14, 15]. To determine more precisely the time course of these synaptotoxic effects, thereby providing a guide to the time points at which to collect samples for phosphoproteomic and transcriptomic analysis, we treated hippocampal cultures for 0.5, 2, 4, and 24 hours with PrP^Sc^ purified from the brains of mice infected with the RML strain of murine-adapted scrapie. Mock-purified material from uninfected brains was used as a negative control. We found that dendritic spine retraction was first detectable after 4 hours of PrP^Sc^ treatment and progressed further by 24 hours (Fig. S1A, B). Since PrP^Sc^ activation of phosphorylation-dependent signaling cascades and subsequent changes in gene expression were likely to lie upstream of, and occur earlier than, observable changes in spine morphology, we decided to include time points earlier than 4 hours in our subsequent experiments. This decision was also motivated by our previous observation that NMDAR-activated Ca^2+^ entry can be detected within 0.5-1 hour, and increased phosphorylation of p38 MAPK by 1 hour following PrP^Sc^ application [15]. Based on these factors, we elected to collect phosphoproteomic data at 1 hour and RNA-seq data at 0.5, 2, 4, and 24 hours after PrP^Sc^ application to hippocampal neuronal cultures (Fig. S1C).

### PrP^Sc^ causes rapid phosphoproteomic alterations associated with synaptic pathways

A key feature of our culture system is that hippocampal neurons are physically separated from the astrocyte feeder layer, allowing collection of proteins and RNA from the neurons with minimal contamination from astrocytes. We undertook an experiment in which five replicate cultures were treated for 1 hour with purified PrP^Sc^, and five cultures for 1 hour with mock-purified material. For proteomic analysis, neuronal protein lysates were processed by a standard workflow that included trypsin digestion, labeling of the resulting peptides with isobaric, tandem mass tags (TMT), isolation of phosphopeptides on Fe-NTA beads, and tandem mass spectrometry (MS/MS) analysis. A portion of each sample prior to phosphopeptide isolation was saved for total proteome analysis. TMT labeling enabled multiplexed, relative quantification of peptide abundances across samples (see PCA plots in Fig. S2A).

In this experiment, we identified 8,217 proteins and 2,135 phosphoproteins with 1,455 overlaps (Fig. S2B). Proteins with │log2fold-change│>0.25 and p-value <0.05 values were considered significant, which led to the identification of 1,212 differentially expressed proteins (62 upregulated and 1,150 downregulated), and 387 differentially phosphorylated proteins (71 increased and 316 decreased) (Fig. S2C, D; Tables S1, S2). To identify pathways associated with the differentially phosphorylated proteins, we performed Gene Set Enrichment Analyses (GSEA) [20] using terms from Gene Ontology (GO) gene sets. Among the most prominently altered pathways (adjusted p<0.05) were those related to cell junctions/projections, axons, synapses, calcium ion binding, and microtubule-based processes (Fig. 1, Table S3). These terms are consistent with the dramatic changes to synaptic structure and function caused by PrP^Sc^, as documented in our previous studies [14, 15].

**Figure 1.**
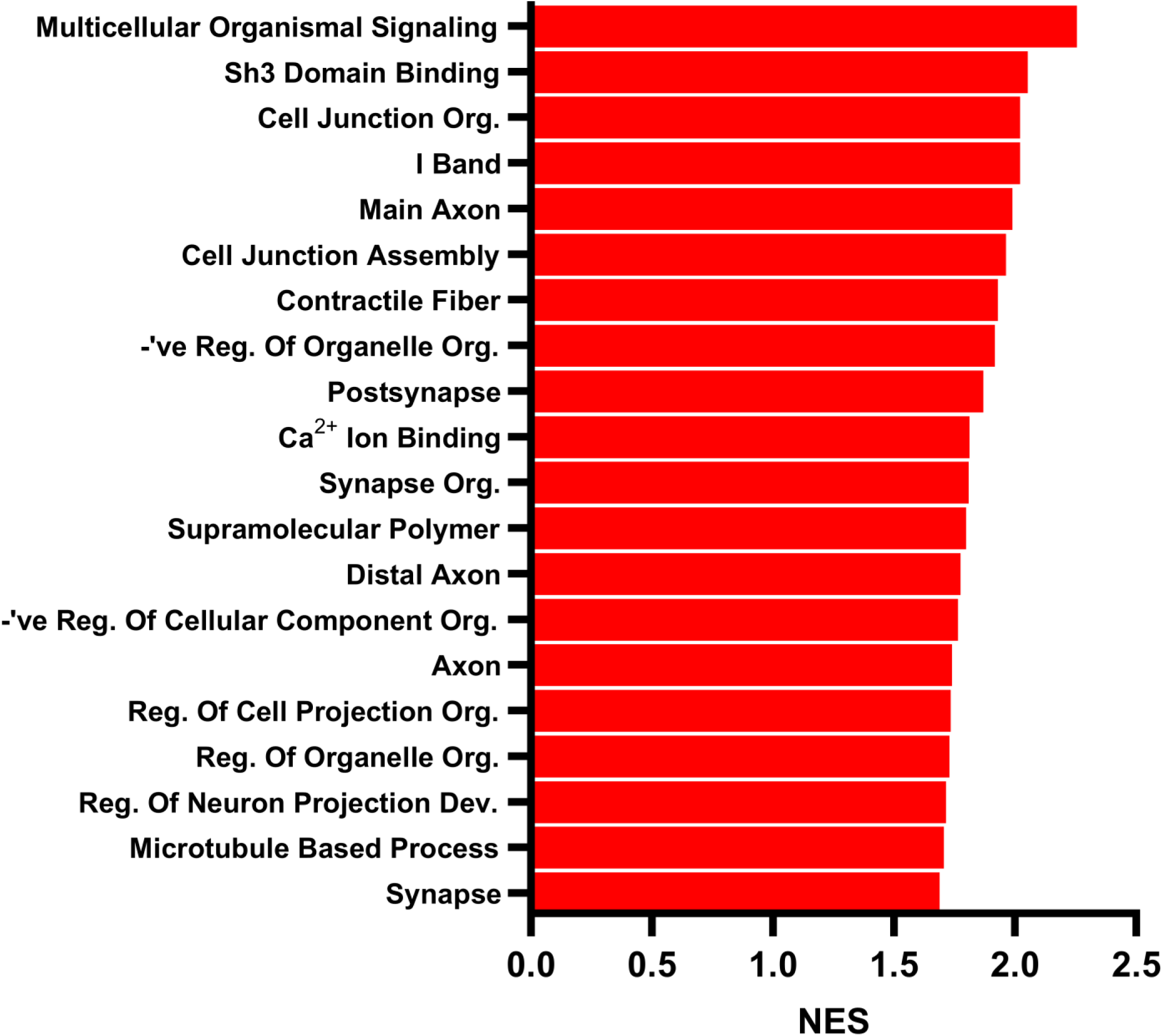
Top pathways altered by PrP^Sc^ treatment of hippocampal neurons based on phosphoproteomic analysis. Pathway enrichment using GO terms for Molecular Function, Biological Process, and Cellular Component was performed on phosphoproteomic data obtained after treatment of neurons with PrP^Sc^ for 1 h. The 20 most highly enriched pathways are ranked by normalized enrichment score (NES), all with adjusted p-values<0.05. See Table S3 for full descriptions of each pathway.

### PrP^Sc^ causes modest transcriptional changes in pathways linked to mitochondria, neurotransmission, and extracellular matrix

Triplicate samples for RNA-seq analysis were collected from hippocampal neurons treated with PrP^Sc^ or mock-purified material for 0.5, 2, 4, and 24 hours. Table S4 lists differentially expressed genes and Fig. S3 shows volcano plots of transcriptional changes at each time point. Using cut-off values for each gene of │log2fold change│>0.25 and p-value <0.05, the number of genes showing significant transcriptional change was modest: 7 upregulated/24 downregulated at 30 minutes; 10 upregulated/22 downregulated at 2 hr; 10 upregulated/12 downregulated at 4 hr; and 86 upregulated/47 downregulated at 24 hrs. Nevertheless, GSEA analysis using the entire transcriptional output yielded statistically significant pathway enrichments (adjusted p-values <0.05) (Fig. 2A-D and Table S5). The most prominent changes were observed at 0.5 hours (upregulation of ribosomal and mitochondrial pathways), likely reflecting an early response to PrP^Sc^-induced oxidative stress, and at 24 hours (upregulation of collagen and extracellular matrix pathways), suggesting remodeling of the extracellular matrix following dendritic spine retraction. The larger number of gene expression changes at 24 hours compared to earlier time points is consistent with a slowly evolving transcriptional response that may occur over several days.

**Figure 2.**
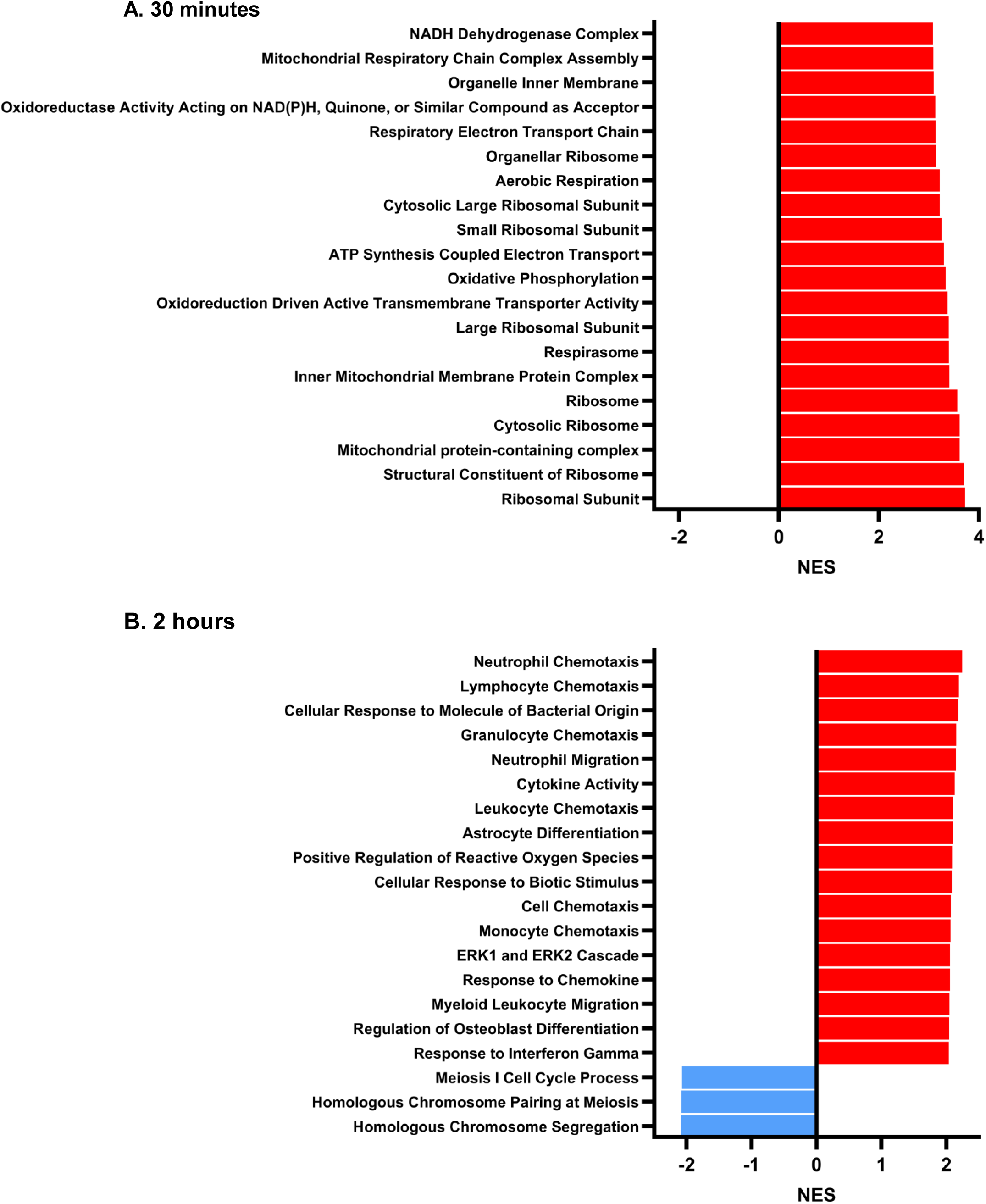
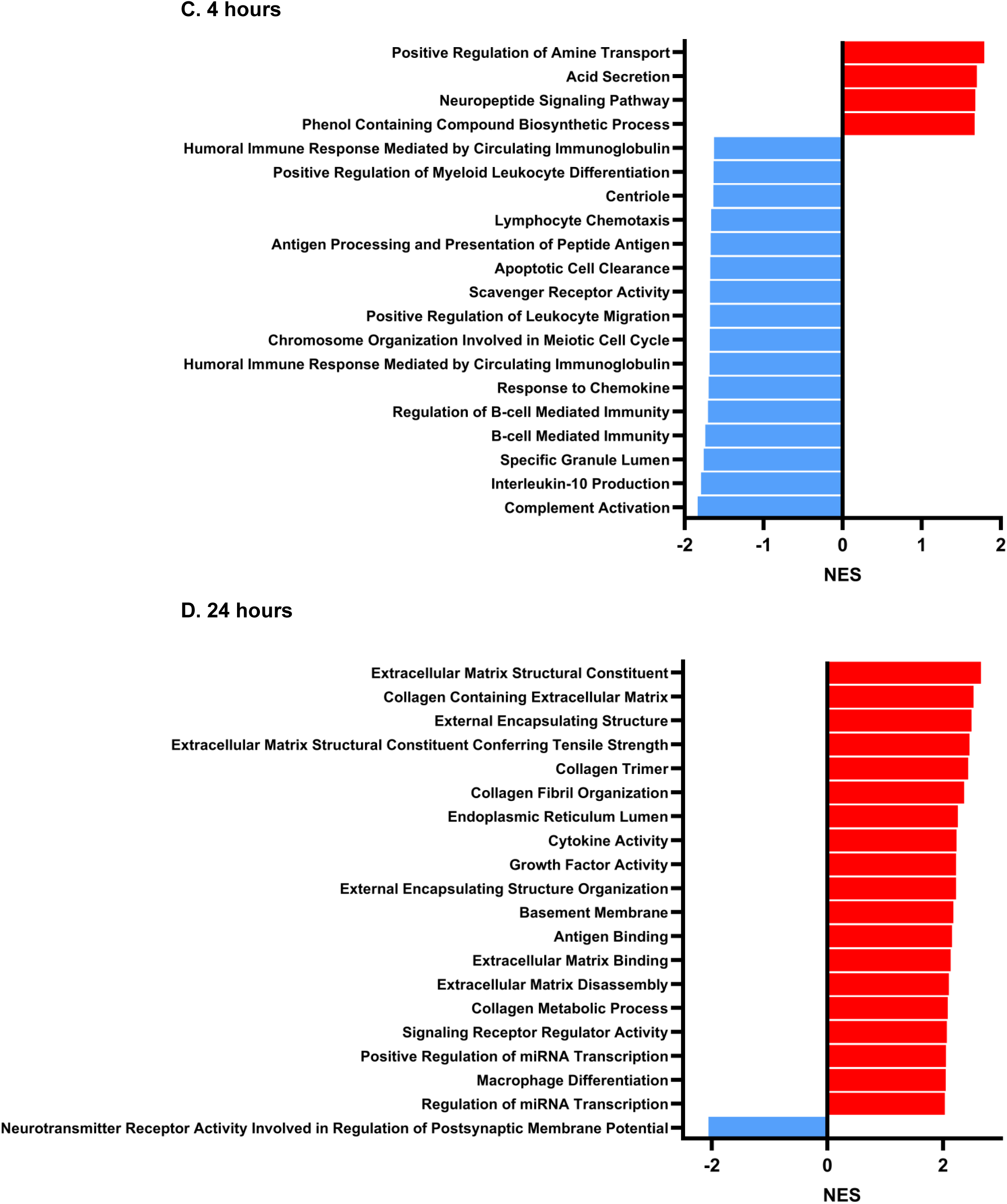
Top pathways altered by PrP^Sc^ treatment of hippocampal neurons based on RNA-seq analysis. GSEA using GO terms for Molecular Function, Biological Process, and Cellular Component was performed on RNA-seq data obtained after treatment of neurons for **(A)** 30 min, **(B)** 2 hrs, **(C)** 4 hrs, and **(D)** 24 hrs. The 20 most highly enriched pathways are ranked by normalized enrichment score (NES). See Table S5 for full descriptions of each pathway.

### Application of a chemo-omic pipeline

We leveraged two public databases, L1000 and P100 [16–19, 21], which contain transcriptomic and phosphoproteomic signatures, respectively, from a panel of cell lines treated with a large number of compounds (referred to as perturbagens). The L1000 database contains over 1 million transcriptomic signatures derived from 9 different core cell lines that have been subjected to >30,000 small molecule perturbagens. The P100 database contains phosphoproteomic signatures from 6 cell lines treated with 90 small molecule perturbagens. Our approach (Fig. 3, and Materials and Methods) was to compare the transcriptomic and phosphoproteomic signatures of PrP^Sc^-treated neurons with the corresponding signatures in these databases, with the goal of identifying compounds that produced the <u>inverse</u> effect on the expression levels and phosphorylation states of each of the input genes or proteins; such compounds would be predicted to reverse the biological effect of PrP^Sc^. The L1000 database was queried using as input the transcriptomic signatures of neurons treated with PrP^Sc^ for 0.5, 2, 4, and 24 hrs, and the P100 database using the phosphoproteomic signature of neurons treated with PrP^Sc^ for 1 hr. Both databases were queried using the CMap portal (https://clue.io/). In the output lists from these queries, we prioritized perturbagens with the most statistically significant anti-correlation with the input signatures, and those with identified molecular targets. In cases where the perturbagens had relatively low specificity for their targets, we identified inhibitors that are more specific, based on literature and commercial sources. The final list of 52 compounds is shown in Table 1.

**Figure 3.**
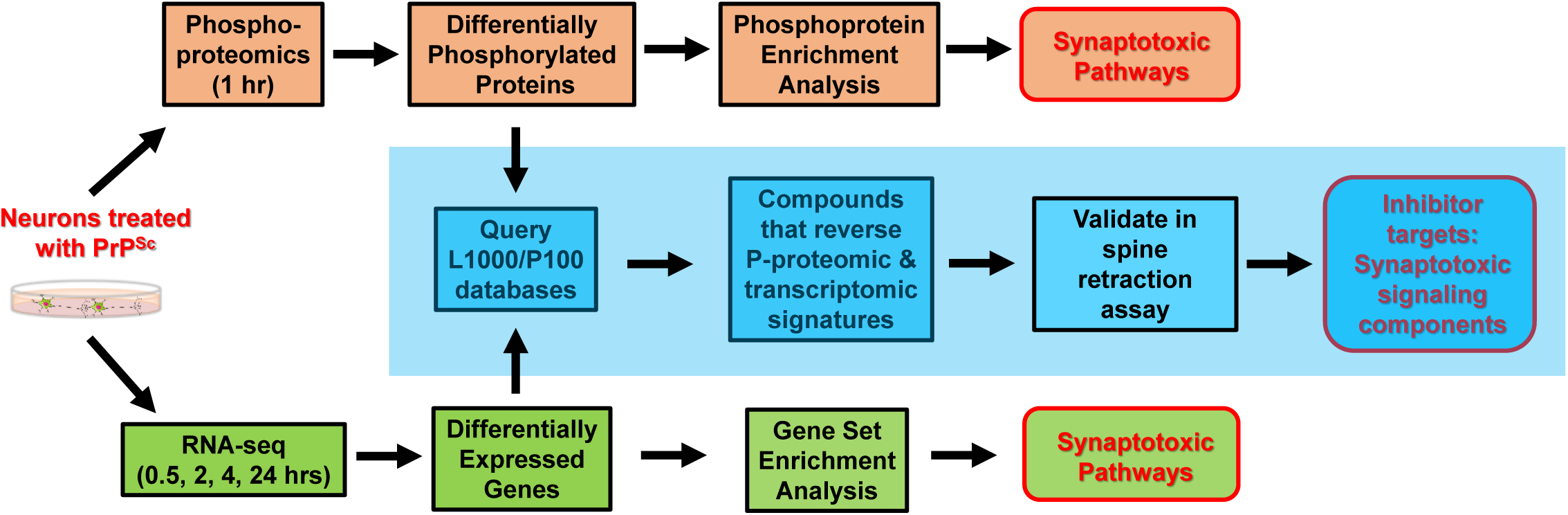
Chemo-transcriptomic-phosphoproteomic pipeline to identify prion synaptotoxic signaling pathways and components. Hippocampal neurons treated with PrP^Sc^ were processed for phosphoproteomics at 1 hr, and for RNA-seq at 30 min, 2 hr, 4 hr, and 24 hr. Synaptotoxic pathways enriched for differentially phosphorylated proteins and differentially expressed genes were identified. Then the lists of differentially phosphorylated proteins and differentially expressed genes were used to interrogate the L1000 and P100 databases, respectively, yielding a final list of 52 candidate compounds that reversed these signatures. These compounds were then tested for their ability to suppress dendritic spine retraction on hippocampal neurons, resulting in a final list of inhibitor targets that represent key components of prion synaptotoxic signaling pathways.

**Table 1.**
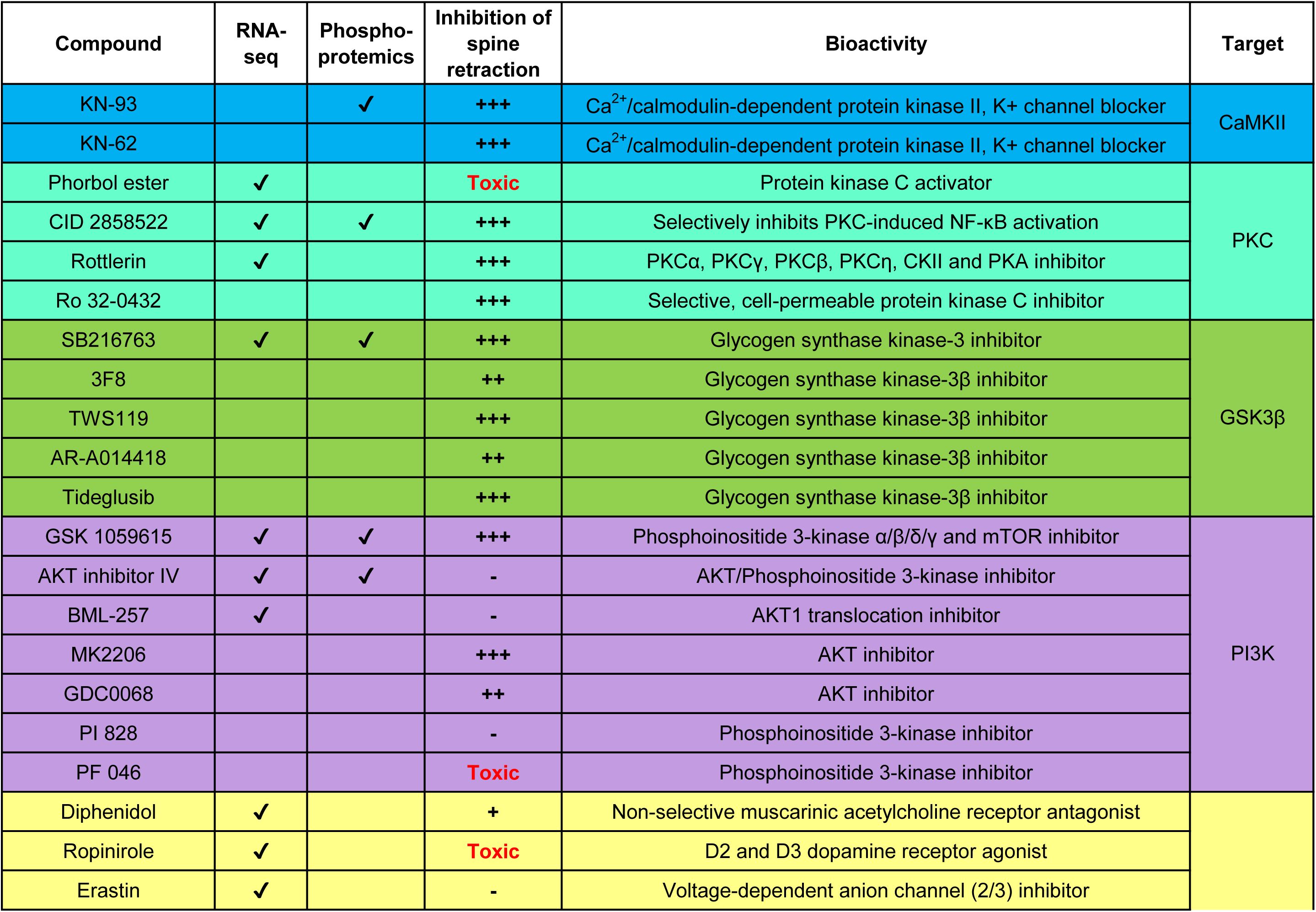
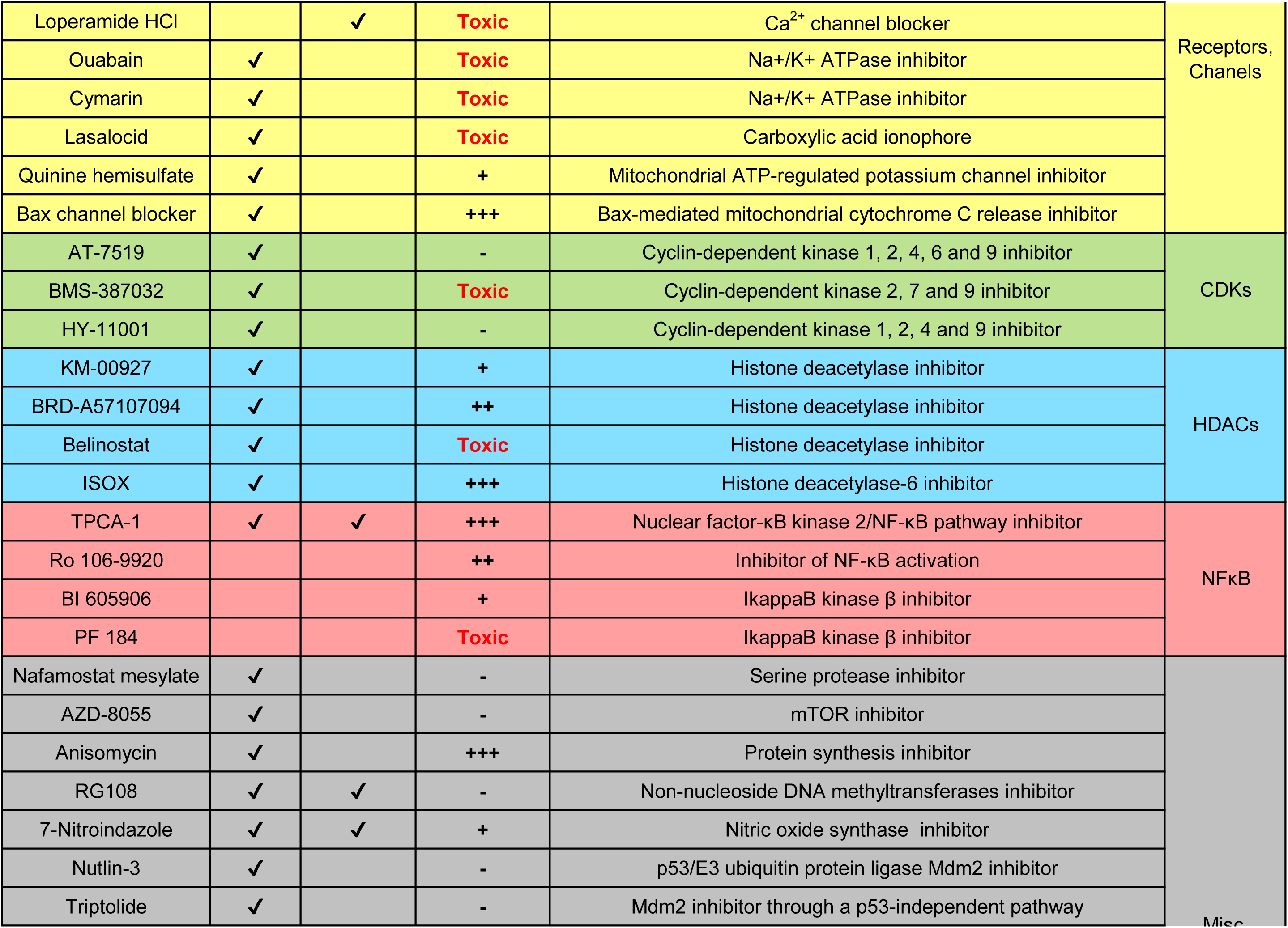
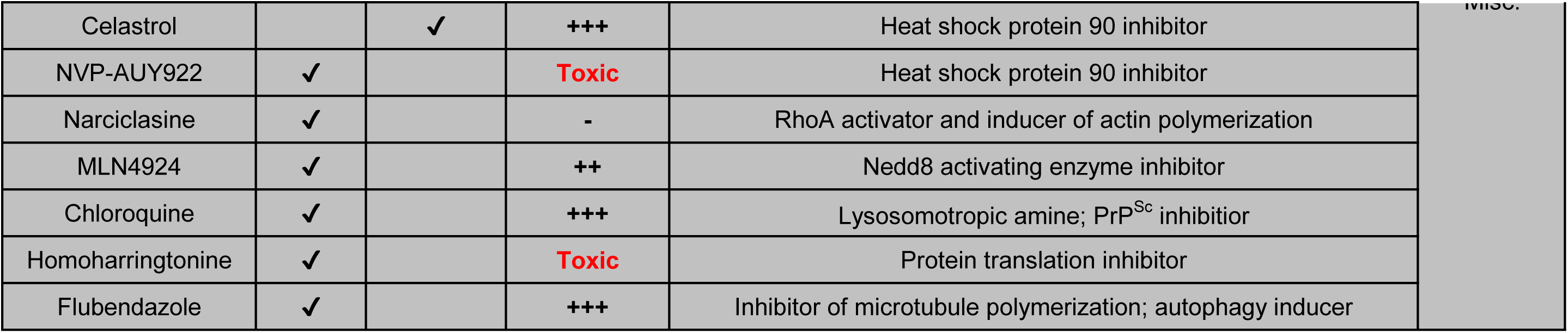
Top-scoring compounds from the chemo-omics pipeline based on their ability to prevent prion synaptotoxicity. Compounds derived from the L1000 transcriptomic databased and the P100 proteomics database are indicated by checkmarks in the columns labeled *RNA-seq* and *Phosphoproteomics*, respectively. Compounds with no checkmarks in either column were selected based on similarity of mechanism of action to compounds derived from the two databases. Compounds are grouped according to their presumed molecular target, shown in the last column of the table, with the three prioritized kinases (CaMKII, PKC, and GSK3β) listed at the top. Dendritic spine retraction was measured with the standard assay [14].. Hippocampal neurons were treated with PrP^Sc^ for 24 hrs at a concentration of 250 nM for each compound. Spine numbers were quantitated after staining with AF488-phalloidin, and inhibition of spine retraction was estimated by comparison to neurons treated with vehicle-only, based on the following scale: +++, maximal effect; ++, moderate effect; +, minimal effect; -, no effect. Compounds that were toxic at 250 nM are indicated.

### A subset of compounds suppresses PrP^Sc^-induced dendritic spine retraction

These 52 compounds were then tested at a concentration of 250 nM for their ability to inhibit dendritic spine retraction in our standard assay, and their effects were scored on a semi-quantitative scale (- to +++) (Table 1 and Fig. S4). Twelve of the compounds were toxic at the tested dose and were eliminated from further consideration. Of the remaining compounds, we identified 17 that maximally suppressed dendritic spine retraction (+++) at a concentration of 250 nM with minimal toxicity. The most consistent effects were observed for inhibitors of three kinases (CaMKII, PKC, and GSK3β), which we decided to prioritize for further analysis by biochemical and cell biological techniques. Fig. 4 shows quantitative measurements of the effects of two different inhibitors of each of these three kinases on dendritic spine number, including appropriate negative controls.

**Figure 4.**
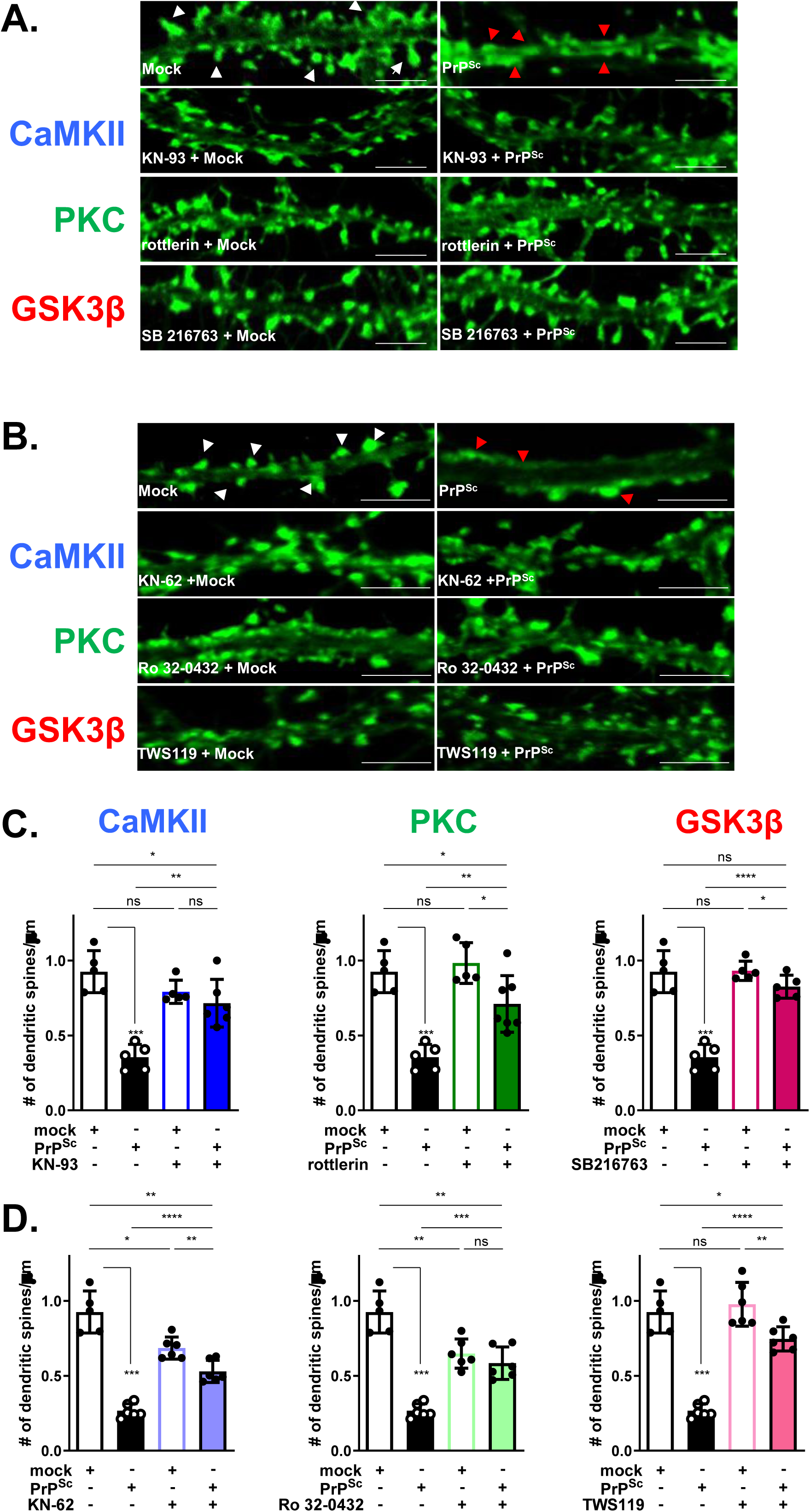
Inhibitors of CaMKII, PKC, and GSK3β prevent spine retraction. **(A,B)** Primary hippocampal neuron cultures were treated with PrP^Sc^ or with mock-purified material (Mock) in the presence of the indicated compounds at 250 nM. Neurons were fixed after 24 hrs and stained with AF488-phalloidin to visualize dendritic spines. In the top row of each panel (no compound), white arrowheads in images of mock-treated neurons and red arrowheads in images of PrP^Sc^-treated neurons indicate healthy and retracted spines, respectively. Scale bars are 5 μm. **(B, C)** Bar graphs show measurements of dendritic spine number per µm for the indicated treatments. Pooled measurements were collected from 5–7 neurons, including 15-25 dendritic regions from at least 2 independent experiments. Data are shown as the mean ± SEM. Statistical analysis was performed using unpaired t-tests; Significance is indicated by ns (not significant), *p < 0.05, **p < 0.01, ***p<0.001, ****P < 0.0001.

CaMKII, PKC and GSK3β are components of well-known and richly connected signaling networks, consistent with the fact that they were all identified based on analysis of both phosphoproteomics and RNA-seq datasets. There is abundant literature documenting their importance in synaptic function (see Discussion). These kinases are themselves regulated by upstream kinases and phosphatases that alter their catalytic activities and subcellular distributions. To further document their contributions to prion synaptotoxic signaling, we therefore wished to investigate changes in their phosphorylation state and intracellular localization in hippocampal neurons exposed to PrP^Sc^.

We initially employed western blotting with phospho-specific antibodies to look for alterations in the phosphorylation states of the three kinases in neuronal lysates but did not detect consistent changes in response to PrP^Sc^ treatment at time points ranging from 1-24 hours (Fig. S5). This result suggested that PrP^Sc^ might be altering the subcellular localization and/or phosphorylation state of only a subset of kinase molecules, thereby changing their distribution with respect to dendritic spines as part of the process of spine collapse, a phenomenon we observed previously for p38 MAPK [15]. We therefore undertook a detailed analysis of the localization of CaMKII, PKC and GSK3β by immunofluorescent staining using antibodies specific for phosphorylated forms of the kinases in conjunction with fluorescent phalloidin to visualize dendritic spines.

### PrP^Sc^ induces rapid translocation of CaMKII to the postsynaptic density

We immunostained hippocampal neurons treated with PrP^Sc^ for 1 hr using antibodies for total CaMKII, two phosphorylated forms of CaMKII (pT286 and pT305/6), and PSD95. Phosphorylation of T286 occurs upon activation of CaMKII, while phosphorylation of T305/6 is inhibitory [22, 23]. We analyzed neurons at 1 hr, well before dendritic spine retraction becomes evident (Fig. S1), in order to capture the earliest events triggered by NMDA-mediated Ca^2+^ influx, which we know occurs within minutes of PrP^Sc^ exposure [15]. We observed a dramatic translocation of total CaMKII to dendritic spines following application of PrP^Sc^ (Fig. 5A, B, E). Interestingly, CaMKII translocation to spines was accompanied by increased staining for PSD95, indicating an initial enlargement of the postsynaptic density (PSD) prior to subsequent spine retraction (Fig. 5C,D, H). When we individually quantitated the amounts of the phosphorylated forms of CaMKII within the PSD, there was a significant decrease in the amount of CaMKII phosphorylated on the inhibitory T305/6 site compared to mock-treated cultures, consistent with preferential translocation of the de-inhibited form of the enzyme (Fig. 5A, B, F). Surprisingly, however, there was a decrease in the proportion of CaMKII-pT286 (Fig. 5C, D, G), the activated form of the enzyme, which may reflect rapid dephosphorylation of translocated molecules at this site by PSD-associated phosphatases. Inhibition of CaMKII with KN-93 prevented translocation of CaMKII to spines, and normalized its ratio with NR2B, the NMDA subunit to which it binds at the PSD when activated (Fig. S6, A, B).

**Figure 5.**
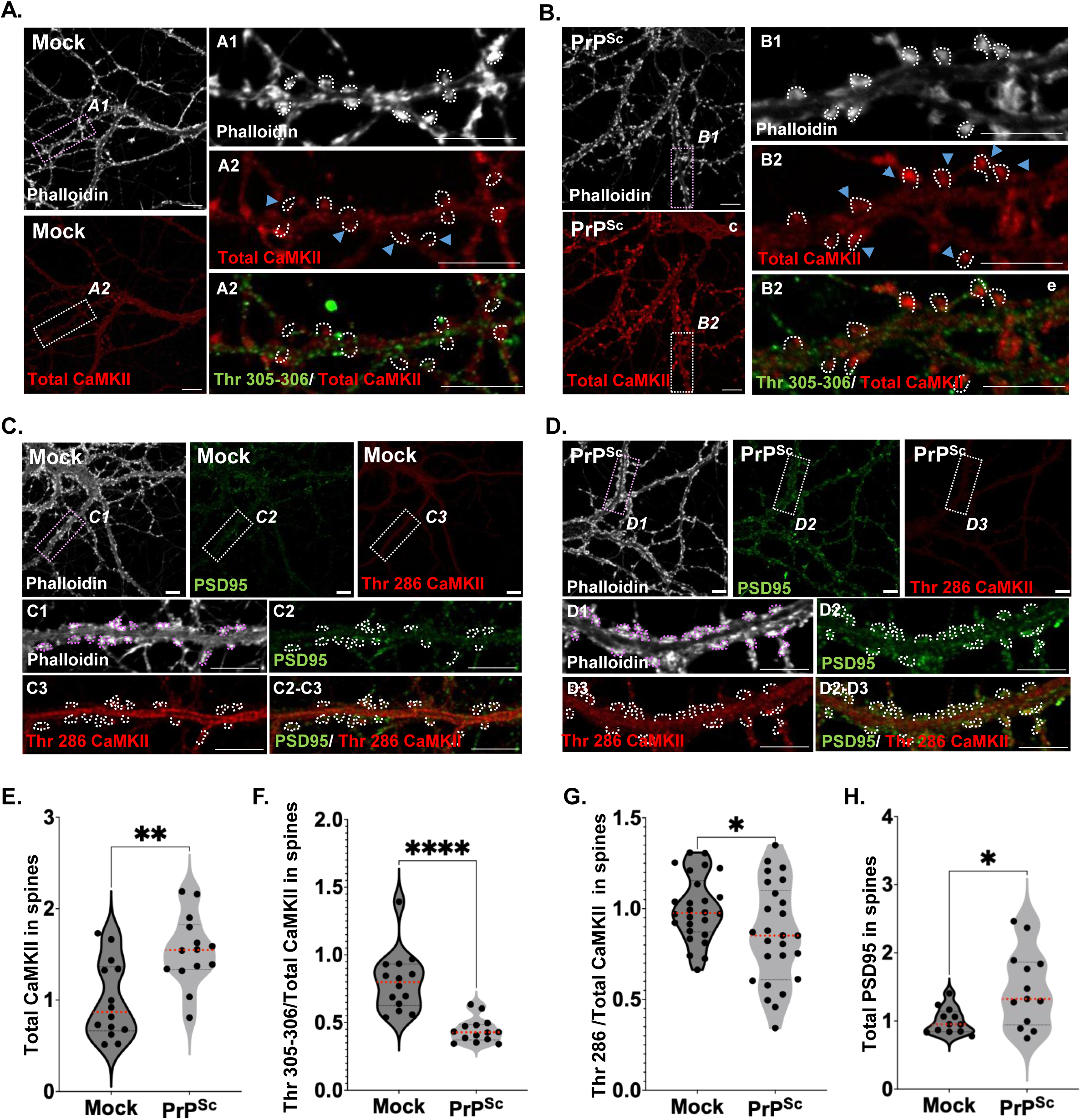
PrP^Sc^ causes rapid translocation of CaMKII to dendritic spines. Hippocampal neurons were treated for 1 hr with mock-purified material **(A, C)** or purified PrP^Sc^**(B, D)**. After fixation, neurons were stained with fluorescent phalloidin (gray), along with antibodies to CaMKII (red) and CaMKII-pT305/306 (green) **(A, B)**; or with antibodies to CaMKII-pT286 (red) and PSD-95 (green) **(C, D)**. Boxed regions in the square panels are shown at higher magnification in the rectangular panels to the right or below (A1-2, B1-2, C1-3, D1-3). Dotted lines in the higher magnification panels outline the positions of intact spines, based on phalloidin staining. Scale bars on all panels are 5 μm. Violin plots show quantitation of Total CaMKII **(E)**, p-Thr305-6/Total CaMKII **(F)**, p-Thr286/Total CaMKII **(G)**, and Total PSD95 **(H)** in spine regions. Measurements were collected from 5-7 neurons, 12-15 dendritic regions from at least 2 independent experiments. Each data point for the PrP^Sc^-treated samples was normalized to the average Mock value from the same experiment. Dotted red lines in each violin plot indicate the median. Statistical analysis was performed on SEM values using unpaired t-tests. Significance is indicated as *p < 0.05, **p < 0.01, ****p < 0.0001.

### PrP^Sc^ causes accumulation of primed PKC in endosomes within the soma and then in dendritic spines

We immunostained hippocampal neurons treated with PrP^Sc^ or mock purified material for 0.5, 2, 4, and 24 hr using antibodies for total PKC and PKC phosphorylated on S660 (Fig. 6). The latter is a priming phosphorylation that is necessary to stabilize newly synthesized molecules of PKC prior to their subsequent activation by signal transduction processes [24]. We found that, at early time points during PrP^Sc^ treatment (0.5 and 2 hrs), PKC-pS660 accumulated prominently in a juxtanuclear compartment within the soma (Fig. 6B, C, G), a pattern similar to what is observed after simulation of neurons with the strong PKC activator phorbol ester (Fig. 6F). This juxtanuclear compartment likely represents a previously described subset of recycling endosomes (the pericentrion), which are thought to amplify the PKC signal during sustained activation [25, 26]. During this same period, there was a decrease in the amount of PKC-pS660 within dendritic spines (Fig. 6B, C, H). By 4 hours, the amount of PKC-pS660 in spines began to increase (Fig. 6D, H), coinciding with significant PrP^Sc^-induced spine retraction observed at this time point (Fig. S1), and suggesting that PKC activation may mediate early synaptic loss. At 24 hours, PKC-pS660 accumulated in both the soma and dendritic spines (Fig. 6E, G, H). Pharmacological inhibition of PKC with Ro 32-0432 prevented these changes in the soma but not in the dendritic spines at 4 and 24 hours (Fig. S7). This timing implicates PrP^Sc^-triggered PKC signaling as a critical mediator of early synaptotoxicity via spine retraction, highlighting 4 hours as a key window for synaptic degeneration driven by PKC-dependent pathways.

**Figure 6.**
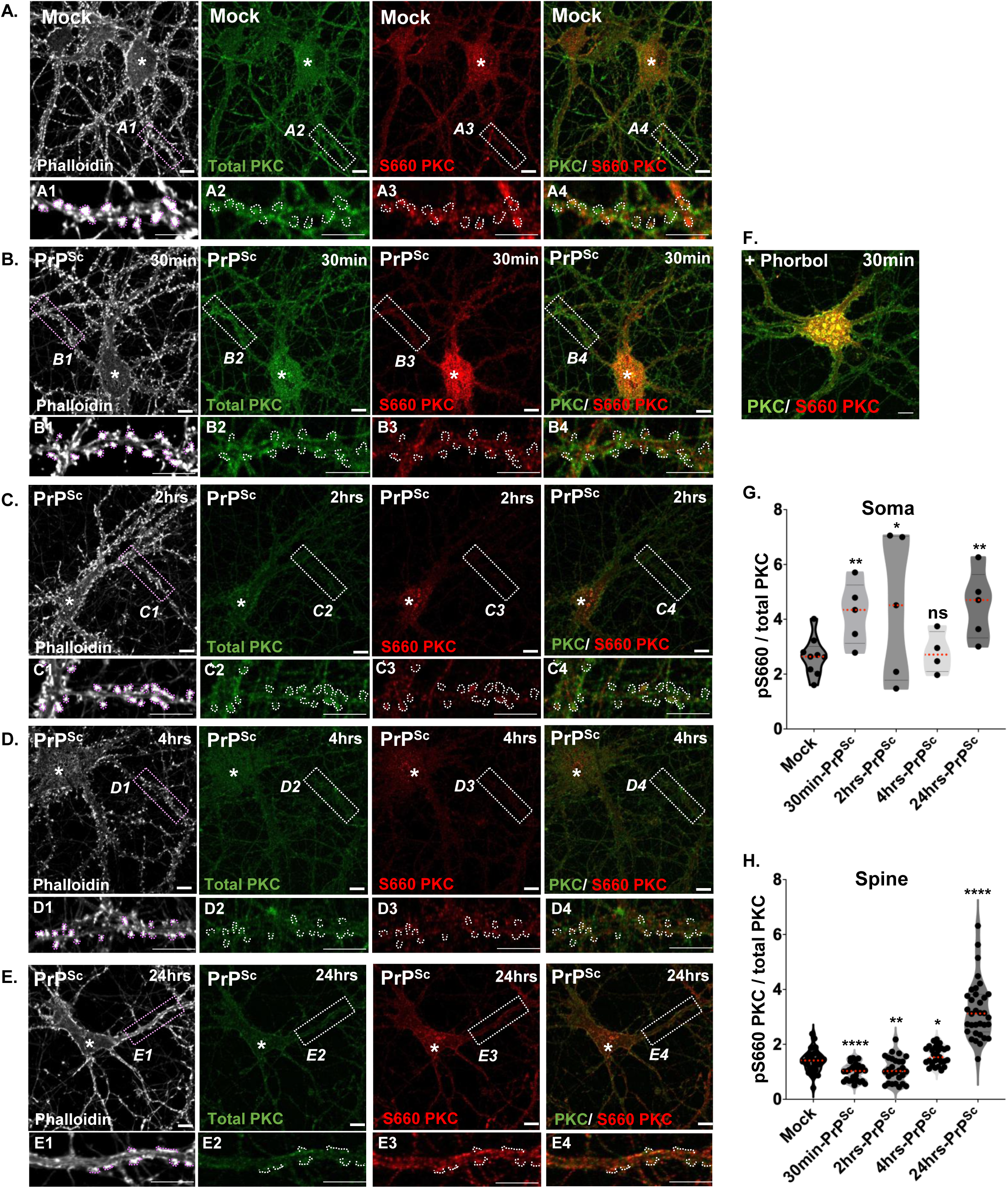
PrP^Sc^ causes accumulation of primed PKC in endosomes within the soma and then in dendritic spines. Hippocampal neurons were treated either with mock-purified material **(A)**, or with purified PrP^Sc^ for 30 min **(B)**, 2 hrs **(C)**, 4 hrs **(D)** or 24 hrs **(E)**. After fixation, neurons were stained with fluorescent phalloidin (gray), along with antibodies to total PKC (green) and PKC-pS660 (red). Boxed regions in the square panels are shown at higher magnification in the rectangular panels below (A1-4, B1-4, C1-4, D1-4). Dotted lines in the higher magnification panels outline the positions of intact spines, based on phalloidin staining. Note the accumulation of vesicles in the soma (indicated by asterisks) that stain for PKC-pS660 after 30 minutes of PrP^Sc^ treatment. This pattern is similar to that seen in neurons treated for 30 minutes with the strong PKC activator phorbol ester **(F)**. Scale bars in all panels = 5 μm. Violin plots show quantitation of PKC-pS660/total PKC ratios within soma regions **(G)** and spine regions **(H)**. Measurements were collected from 5-7 neurons, 30-45 dendritic and somatic regions from at least 2 independent experiments. Each data point for the PrP^Sc^-treated samples was normalized to the average Mock value from the same experiment. Dotted red lines in each violin plot indicate the median. Statistical analysis was performed on SEM values using unpaired t-tests. Significance is indicated as ns (not significant), *p < 0.05, **p < 0.01, ****p < 0.0001.

### PrP^Sc^ causes accumulation of activated GSK3**β** within dendritic spines

We expected that changes in the phosphorylation state of GSK3β would occur by auto-phosphorylation of the Y216 activating site, since GSK3β inhibitors prevented dendritic spine retraction after 24 hrs of PrP^Sc^ treatment (Table 1, Figs. 4 and S4). To test this prediction, we stained hippocampal cultures treated with PrP^Sc^ for 1 hr with antibodies to GSK3β-pY216 and total GSK3β and then imaged the ratio of the two signals in the region of dendritic spines, as marked by phalloidin staining. As predicted, we found that the amount of GSK3β-pY216 in dendritic spines was significantly increased after 1 hour of PrP^Sc^ treatment (Fig. 7A, B). Two GSK3β inhibitors (Tideglusib and AR-A014418) were able to suppress PrP^Sc^-induced spine retraction with K_i_ values of ∼30 nM (Fig. 7C-E).

**Figure 7.**
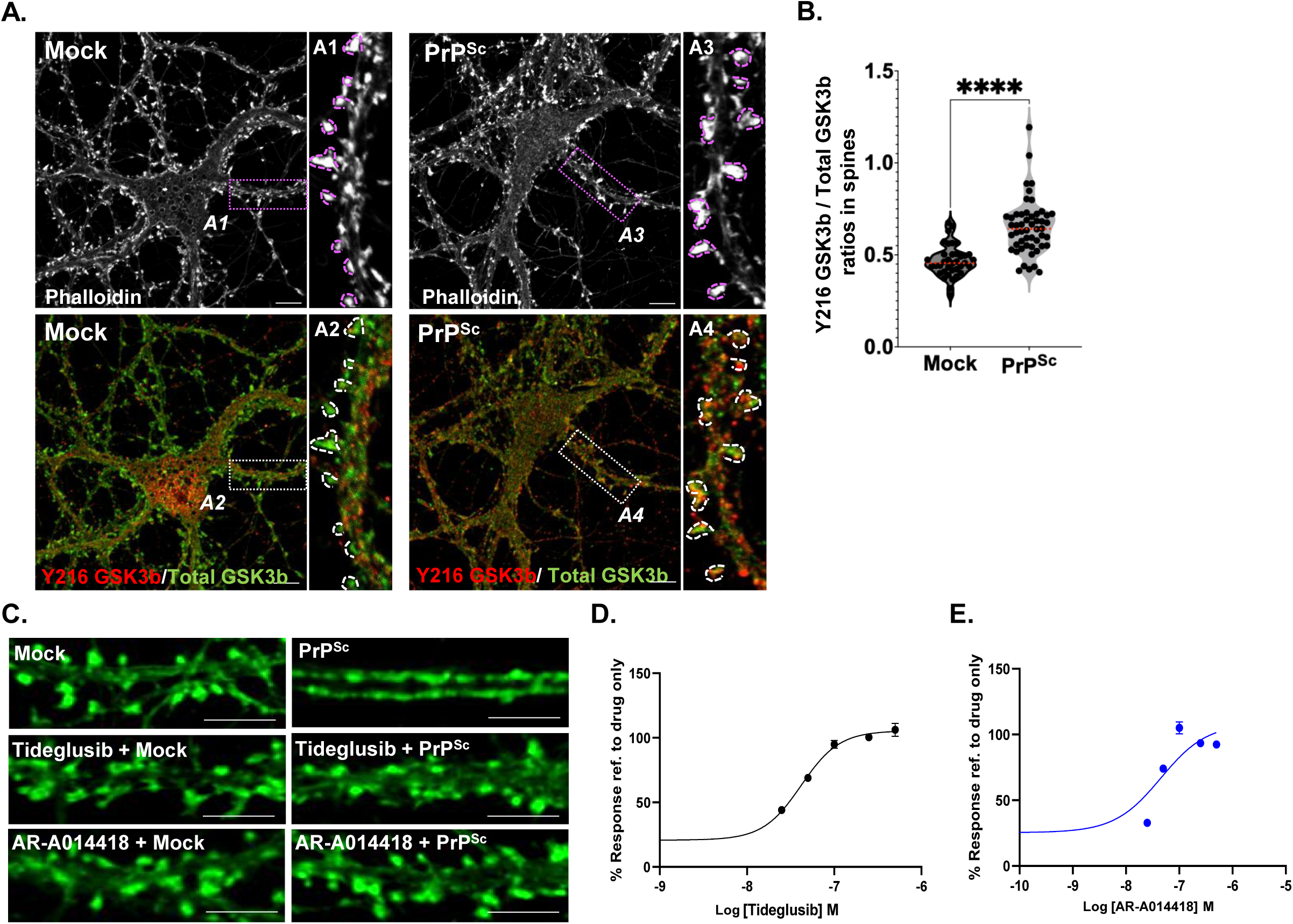
PrP^Sc^ increases active pY216-GSK3β in dendritic spines, and this effect is prevented dose-dependently by GSK3β inhibitors. **(A)** Hippocampal neurons were treated for 1 hr with mock-purified material (Mock, left-hand group of panels) or purified PrP^Sc^ (PrP^Sc^, right-hand group of panels). After fixation, neurons were stained with fluorescent phalloidin, along with antibodies to total GSK3β and GSK3β-pY216. Upper panels show images of phalloidin staining (gray). Lower panels show merged images of total GSK3β (green) and GSK3β-pY216 (red) staining. The boxed regions in the large, square panels are shown at higher magnification in smaller, rectangular panels to the right (A1-4). Dotted lines in the smaller panels outline the positions of intact spines, based on phalloidin staining. Spines in PrP^Sc^- treated neurons (lower right) show enhanced GSK3β-pY216 /total GSK3b staining (yellow) compared to mock (lower left). **(B)** Violin plot showing quantification of GSK3β-pY216/total GSK3β within spine regions, normalized as described in Figs. 5 and 6. Significance is indicated as ****p<0.0001. **(C)** Representative fluorescence micrographs showing inhibition of spine retraction by Tideglusib and AR-A014418 (both at 30 nM). Neurons were pretreated for 2 hrs with the compounds, after which they were treated in the continued presence of the drugs for 24 hrs with either mock-purified material or purified PrP^Sc^. After fixation, neurons were stained with Alexa 488-labeled phalloidin for visualization and counting of dendritic spines. The top pair of panels show neurons that were not pretreated with inhibitors. Scale bars = 5 μm. **(D, E)** Dose response curves for inhibition of PrP^Sc^-induced dendritic spine retraction by GSK3β inhibitors. Conditions were the same as in panel C. Pooled measurements were collected from 30-45 dendritic regions on 5–7 neurons from at least 2 independent experiments. The y-axis shows spine density (number/µm) in PrP^Sc^-treated neurons exposed to drug as a percentage of spine density in mock-treated neurons exposed to drug. Both drugs inhibited spine retraction with K_i_ values of ∼30 nM.

## DISCUSSION

In this study, we have used a chemo-omic pipeline to identify three kinases (CaMKII, PKC, and GSK3β) that play key roles in the earliest events in prion synaptotoxic signaling. The pipeline used as input phosphoproteomic and transcriptomic data from hippocampal neurons treated with PrP^Sc^, which was then compared to databases of cellular responses to millions of small molecules. The resulting output converged on a set of inhibitors of the three kinases that prevented PrP^Sc^-induced dendritic spine retraction in cultured hippocampal neurons. Finally, we confirmed a role for these kinases in synaptic degeneration by showing that the active forms of the kinases translocated to dendritic spines as soon as one hour after PrP^Sc^ application. Taken together, our results define the earliest steps in a prion synaptotoxic signaling cascade, and they identify a host of new inhibitors and potential drug targets for therapeutic purposes.

### Pathway analysis

Prior to applying the chemo-omic pipeline, we performed gene set enrichment analysis (GSEA) of the phosphoproteomic and transcriptomic data to identify cellular pathways altered by PrP^Sc^ treatment of cultured hippocampal neurons. Significant changes in the phosphoproteome were detected after 1 hr of PrP^Sc^ treatment, consistent with the hypothesis that acute exposure of neurons to prions activates a network of phosphorylation-dependent signaling pathways that are triggered by the initial interaction of PrP^Sc^ with cell-surface PrP^C^ [13, 15]. Gene set enrichment analysis of the phosphoproteomic data returned GO terms related to cell junctions/projections, axons, synapses, calcium ion binding, and microtubules among the top 20 most highly regulated pathways. A role for phosphorylation-dependent processes in prion synaptotoxicity is not surprising, since specific kinases and phosphatases are known to be involved in remodeling of dendritic spine morphology and changes in synaptic transmission [27].

In contrast to the phosphoproteomic changes, the transcriptomic changes we observed in PrP^Sc^-treated neurons were relatively modest across all time points (<50 genes up or down regulated). Nevertheless, GSEA analysis using the entire transcriptome yielded statistically significant pathway enrichments (adjusted p-values<0.05). The most prominent enrichments were observed at 0.5 hours (upregulation of ribosomal and mitochondrial pathways), likely reflecting an early response to PrP^Sc^-induced oxidative stress, and at 24 hours (upregulation of collagen and extracellular matrix pathways), indicative of synaptic degeneration and consequent remodeling of the ECM. The modest effect of PrP^Sc^ on neuronal gene expression likely reflects the fact that, even after 24 hours of PrP^Sc^ treatment, when substantial numbers of dendritic spines have retracted and synaptic transmission has declined, neuronal viability is unaffected and dendritic shafts remain intact. The pattern of gene expression that we observe here is consistent with RNA-seq studies of the brains of prion-infected mice, in which neurons display relatively few transcriptional changes until the end-stage of the disease, with the earliest detectable changes occurring in astrocytes and microglia during the presymptomatic phase [28–30]. Taken together, our results suggest that the earliest response of neurons to prion exposure occurs at the level of protein phosphorylation-dependent signaling, prior to major compensatory changes in gene expression.

### Chemo-omic pipeline

A novel aspect of this study is the use of a bioinformatic pipeline in which we used our transcriptomic and phosphoproteomic data to interrogate a set of public databases (L1000, P100) containing transcriptomic and phosphoproteomic signatures of a standard set of cell lines treated with large numbers of compounds at different concentrations for varying lengths of time [16–19]. The objective of this procedure was to identify compounds that produced signatures that were the <u>inverse</u> of the input signatures derived from PrP^Sc^-treated hippocampal neurons. We predicted that such compounds would reverse the biological effect of PrP^Sc^, as assayed by measurements of dendritic spine retraction. This approach was first described by Lamb *et al*. [17], and there are now several published precedents for its successful application in disease contexts, including the identification of drugs for treatment of AD [31, 32], atherosclerosis [33], SARS-CoV-2 infection [34], and cancer [35]. All of these cited studies used transcriptomic data to query the L1000 database; to our knowledge, ours is the first to interrogate the L1000 and P100 databases using both transcriptomic and phosphoproteomic data, respectively, and to integrate the output from both sources.

### CaMKII

CaMKII has long been associated with synaptic changes underlying learning, memory, and neurodegenerative conditions [22, 23]. The participation of CaMKII in prion synaptotoxicity is consistent with the rapid influx of Ca^2+^ into cultured hippocampal neurons that we observe within minutes of exposure to PrP^Sc^, which would increase binding of Ca^2+^ to calmodulin (CaM), causing activation of CaMKII [15]. This Ca^2+^ influx is mediated primarily by NMDARs, since it is abolished by NDMAR blockers, which also prevent dendritic spine retraction. Canonically, during the process of long-term potentiation (LTP), Ca^2+^-CaM binding to CaMKII induces phosphorylation of T286, resulting in autonomous kinase activation, and translocation of CaMKII to the PSD, where it binds to the C-terminal cytoplasmic tail of NR2B-containing NMDA receptors [22, 23]. Consistent with this model, we observed a large shift of CaMKII to the dendritic spines of hippocampal neurons exposed to PrP^Sc^ (Fig. 5). Interestingly, however, the PSD-localized CaMKII is not phosphorylated at T286, perhaps reflecting the action of phosphatases that rapidly dephosphorylate the active enzyme upon arrival at the PSD. In contrast, spine-localized CaMKII showed reduced phosphorylation at T305/6, an inhibitory site, consistent with prior activation of the kinase. The initial burst of CaMKII translocation to dendritic spines, coupled with the transient increase in PSD95 staining we observed (Fig. 5), is consistent with a transient enlargement of spines, which is then followed by spine collapse at later times. Activated CaMKII has been shown to regulate the actin cytoskeleton in spines via Ca^2+^-induced phosphorylation of cytoplasmic actin-binding proteins [36], a process that likely contributes to the enlargement and then retraction of spines in response to PrP^Sc^. Treatment with the CaMKII inhibitor KN-93 reversed the initial translocation of CaMKII to dendritic spines, normalized the CaMKII/NR2B ratio, and ultimately prevented spine collapse following PrP^Sc^ treatment (Figs. 4 and S6). It remains to be determined whether PrP^Sc^-dependent activation of NMDARs is due to direct binding of PrP^Sc^ to NMDAR subunits, binding other membrane proteins, or perturbation of the lipid bilayer [13, 37].

### PKC

We found that PrP^Sc^ causes rapid accumulation of primed PKC in endosomes within the soma, followed by its translocation to dendritic spines. PKC inhibitors prevent these effects, and block PrP^Sc^-induced dendritic spine retraction. PKC is known to play multiple roles at the synapse, including mediating structural changes accompanying synaptic plasticity, enhancing synaptogenesis, modulating Ca^2+^ currents, and integrating signal transduction pathways [24]. Of particular relevance here, PKCα is activated in response to NMDAR-mediated Ca^2+^ influx during LTP, causing translocation of the kinase to synapses where it binds to PSD-95 and other PSD scaffolding proteins via its PDZ-binding domain, further potentiating NMDAR signaling [24, 38, 39]. It is likely that the large NMDAR-mediated Ca^2+^ influx elicited by PrP^Sc^ [15] pathologically stimulates this same pathway. Of note, we find that PKC translocated to dendritic spines following PrP^Sc^ stimulation is phosphorylated on residue S660. This site is one of three priming sites that become auto-phosphorylated upon synthesis of the PKC protein, a process that is necessary to stabilize the nascent polypeptide and prevent its degradation [24, 40]. We find that the PrP^Sc^-induced accumulation of primed PKC in endosomal compartments within the soma mimics the effect of the strong PKC activator, phorbol ester (Fig. 6F). These endosomal compartments likely correspond to the pericentrion, a subset of juxtanuclear, recycling endosomes seen in non-neuronal cells treated with phorbol ester [26, 41]. Taken together, our observations demonstrate that prions produce a powerful and sustained activation of the PKC pathway.

Class 1 metabotropic glutamate receptors (mGluR1 and mGluR5), which are coupled to Gα_q_, are canonical upstream activators of PKC via their stimulation of phospholipase C, which generates diacylglycerol and IP3 second messengers [24]. However, we have found that dendritic spine retraction in hippocampal neurons in response to PrP^Sc^ is not blocked by the mGluR5 inhibitors MPEP and CTEP [15, 42], suggesting that this receptor is not involved in PKC-mediated spine retraction, and that NMDAR-mediated Ca^2+^ influx may play a more important role in acutely activating the PKC pathway. High concentrations of Ca^2+^ can mediate binding of PKC to anionic membrane lipids in the absence of diacylglycerol, and this can lead to phosphorylation of PSD95 [43]. However, mGluR1 and mGluR5 may contribute at later stages of disease, as suggested by the observation that inhibitors of these receptors block neuronal loss in brain slice cultures infected with prions or treated by prion-mimetic antibodies, and also mitigate clinical symptoms and neuropathology in prion-inoculated mice [42, 44]. Thus, these receptors may be important for degenerative changes downstream of acute synaptotoxicity. mGluR5 has also been shown to function as a co-receptor with PrP^C^ to bind Aβ oligomers in Alzheimer’s disease [45]. Since mGluR5 inhibitors block retraction of dendritic spines on cultured hippocampal neurons exposed to Aβ oligomers but not PrP^Sc^, it is likely that these two oligomers acutely activate distinct synaptotoxic pathways [15].

### GSK3**β**

We have shown that GSK3β inhibitors block PrP^Sc^-induced dendritic spine loss, and that PrP^Sc^ treatment causes accumulation within spines of the constitutively active form of GSK3β that is phosphorylated on residue Y216. GSK3β is a multi-functional Ser/Thr kinase that is enriched in dendritic spines, and is involved in both physiological and disease processes in the brain via phosphorylation of a number of key substrates, including PSD95, KLC3 and tau [46, 47]. Unlike many other kinases, GSK3β is constitutively active due to autophosphorylation at Y216, and upstream regulators typically act to reduce GSK3β enzymatic activity via an inhibitory phosphorylation at residue S9 [48]. Relevant to the results presented here, GSK3β plays an important role in the induction of NMDA receptor-dependent long-term depression (LTD) during which GSK3β translocates to the PSD and phosphorylates PSD95, leading to AMPAR internalization [49]. Moreover, phosphorylation of phosphatidylinositol 4 kinase type IIα (PI4KIIα) by GSK3β stabilizes a constitutive pool of NMDARs at the synapse [46, 50]. Over-activation of the latter pathway by PrP^Sc^ could therefore increase the number of synaptic NMDARs, thereby augmenting Ca^2+^ influx and activation of CaMKII.

Interestingly, GSK3β also plays a key role in Alzheimer’s disease, primarily by its ability to phosphorylate the microtubule-associated protein, tau [47]. In this context, Aβ oligomers activate GSK3β, possibly via the kinase Pyk, leading in turn to enhanced phosphorylation of tau and its aggregation into amyloid fibrils [51–53]. GSK3β inhibitors reverse electrophysiological and behavioral deficits in AD mouse models by targeting this pathway, and they have been tested in clinical trials for treatment of AD [54]. Although upstream components of PrP^Sc^ and Aβ synaptotoxic pathways are distinct (see above), it is possible that they share downstream, GSK3β-dependent processes.

### A kinase-dependent signaling network for prion synaptotoxicity

Collectively, our results provide evidence for a signal transduction network stimulated by PrP^Sc^ that contains four major nodes: NMDARs, CaMKII, PKC, and GSK3β (Fig. 8). In this scheme, PrP^Sc^ generated on the cell surface activates NMDARs, leading to massive Ca^2+^ influx, phosphorylation of CaMKII and PKC, and translocation of these kinases to the PSD, where they further potentiate NMDAR channel activity and trafficking. GSK3β may augment these responses by stabilizing NMDARs at the synapse. Importantly, this signaling network utilizes the same molecular components that are responsible for LTP and LTD, suggesting that prions co-opt normal, physiological processes underlying learning and memory to produce neurotoxic effects. We have recently shown that newly converted, membrane-anchored PrP^Sc^, rather than exogenous, extracellular PrP^Sc^, is the trigger for prion synaptotoxicity, suggesting that conversion of PrP^C^ to PrP^Sc^ on the plasma membrane flips the downstream signaling network into an over-drive mode [13]. It remains to be determined whether PrP^Sc^-dependent activation of NMDARs is due to direct binding of membrane-anchored PrP^Sc^ to NMDAR subunits, indirect interaction via other membrane proteins, or perturbation of the lipid bilayer [37].

**Figure 8.**
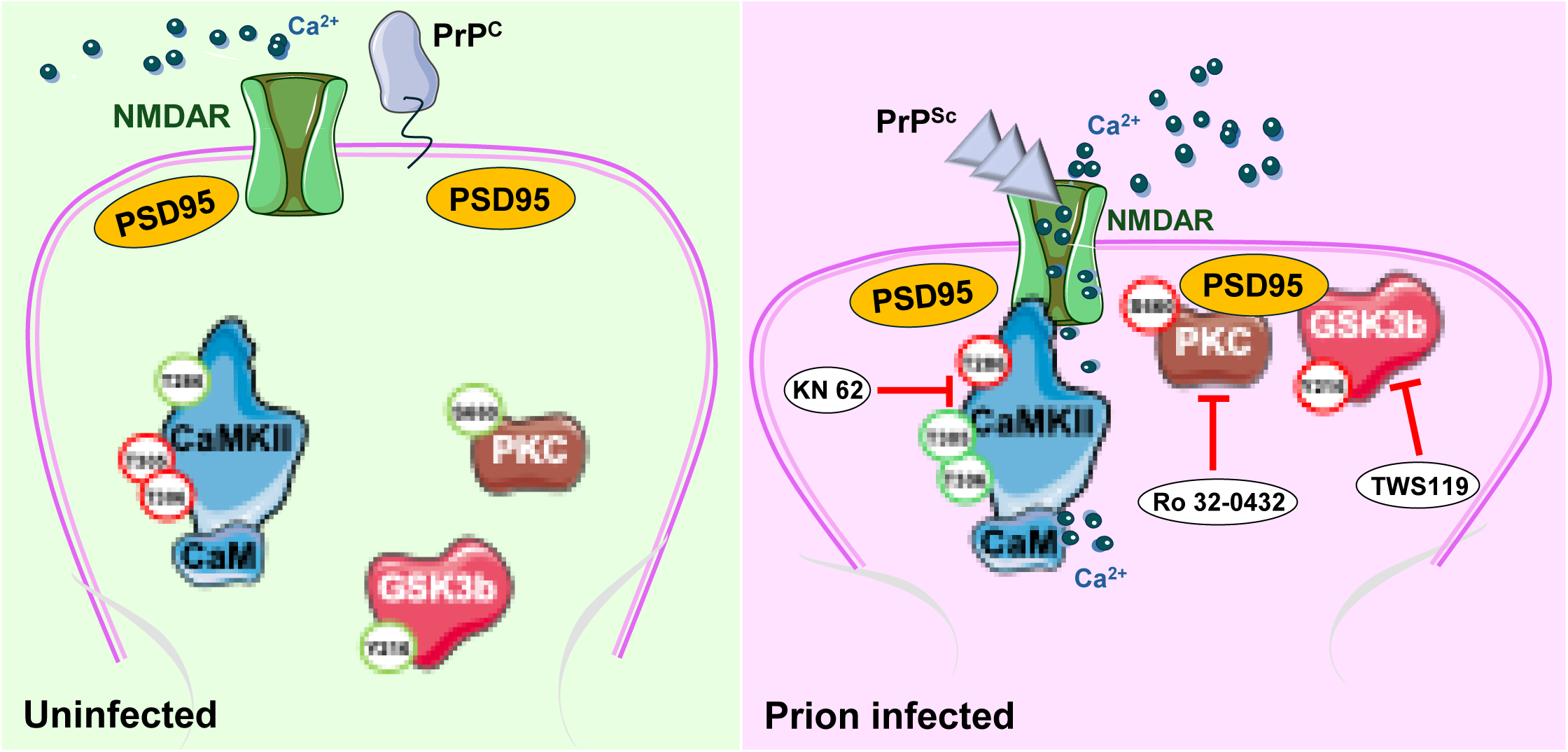
Schematic showing effect of PrP^Sc^ on CaMKII, PKC, and GSK3β in dendritic spines. In the initial stage of prion infection, PrP^Sc^ activates NMDARs in the postsynaptic membrane, either directly as shown here, or indirectly via interaction with other membrane proteins or the lipid bilayer (not shown). The resulting influx of Ca^2+^ ions activates CaMKII by phosphorylation of T286 and dephosphorylation of T305/306, accompanied by CaMKII translocation to the PSD where it binds to NMDARs via the cytoplasmic tails of NR2B subunits (not shown). PrP^Sc^ also causes primed PKC phosphorylated on S660, as well as activated GSK3β phosphorylated on Y216, to translocate to the PSD, where they can associate with PSD-95. Together, these signaling events lead to collapse of the actin cytoskeleton and retraction of the dendritic spine. Inhibitors of CaMKII, PKC, and GSK3β (KN-62, Ro 32-0432, and TWS119, respectively) prevent spine retraction. Amino acid sites in the three kinases that are phosphorylated are shown in red circles, and those that are dephosphorylated are shown in green circles.

In a previous study [15] using the same neuronal culture system as employed here, we identified p38 MAPK as a key player in prion-induced synaptotoxicity, based on both pharmacologic and genetic evidence. PrP^Sc^ treatment of hippocampal neurons increased the amount of phosphorylated p38 MAPK in dendritic spines, and specific p38 MAPK inhibitors, as well as a dominant negative mutant of p38 MAPK, prevented PrP^Sc^-induced retraction of spines and associated decrements in synaptic transmission. Although p38 MAPK was not prioritized in the chemo-omic pipeline used here, there is ample evidence connecting it to the kinase-dependent signaling network proposed above. p38 MAPK directly phosphorylates GSK3β on serine 389/390, possibly modulating its susceptibility to other regulatory phosphorylations [55]. p38 MAPK is also activated during NMDAR-dependent LTD and contributes to NMDAR/Ca^2+^-mediated excitotoxic damage [56, 57].

Several other studies have highlighted a role for Ca^2+^-mediated excitotoxic processes in prion neurotoxicity. For example, it has been reported that prion infection results in upregulation of *Arc/Arg3.1*, an immediate early response gene activated by neuronal activity [58, 59]. Our observation that PrP^Sc^ stimulates rapid, NMDAR-dependent Ca^2+^ influx [15] followed by activation of Ca^2+^-dependent signaling processes (this work) is consistent with this idea. Although there have not been previous global, phosphoproteomic analyses of prion infected mice or cultured cells, a relevant recent study showed that expression of a neurotoxic mutant PrP in mouse brain or cultured neurons enhances NMDAR activity, activates CaMKII and PKC, and triggers excitotoxic damage [60]. A similar mechanism may be operative in mice and cells we created that express a different neurotoxic PrP mutant, which is associated with aberrant ion channel activity [61–63]. These mutant PrP molecules may be triggering some of the excitotoxic signaling processes that are induced by prion infection.

Our results highlight possible therapeutic avenues for treatment of prion diseases. Indeed, the chemo-transcriptomic-phosphoproteomic pipeline we have employed here has typically been leveraged to identify and repurpose therapeutic compounds [31, 33–35]. Inhibitors of NMDARs, CaMKII, PKC, GSK3β, and p38 MAPK have all been tested for treatment of AD and neurological disorders [64], but only memantine, an NMDAR open-channel blocker, has been tested in prion-infected mice, where it was shown to extend survival times [65]. Our results suggest that further investigation of these inhibitors as treatments for prion diseases, particularly in the presymptomatic phase, is warranted. These compounds, along with others we have discovered using cell-based screening assays [66–69], may represent useful adjuncts to treatments based on reducing PrP^C^ expression [70].

## MATERIALS AND METHODS

### PrP^Sc^ purification

PrP^Sc^ purification from the brains of terminally ill, RML prion-inoculated C57BL/6J mice was performed as described previously utilizing pronase digestion and sodium phosphotungstate precipitation [71] Parallel, mock purifications were performed using age-matched, uninfected brains. Purity was assessed by SDS-PAGE with silver staining and western blotting. In all experiments, purified PrP^Sc^ was added to neuronal cultures at a final concentration of 4.4 μg/ml to be consistent with our previous studies [14, 15]. Mock-treated samples received an equivalent amount of material purified from uninfected brain tissue.

### Primary hippocampal neuronal cultures

Timed-pregnant C57BL/6J mice (wild-type, WT) were obtained from The Jackson Laboratory (Bar Harbor, ME). All procedures involving animals were conducted in accordance with the U.S. Department of Agriculture Animal Welfare Act and the NIH Policy on Humane Care and Use of Laboratory Animals. Hippocampal neurons were isolated from postnatal day 0 (P0) pups as described previously [72]. Two types of neuronal-astrocyte culture systems were used.

For dendritic spine analysis, neurons were plated at low density (75 cells/mm²) on poly-L-lysine-coated coverslips. After several hours, coverslips were inverted over an astrocyte feeder layer, with each coverslip held just above the astrocyte layer by several wax dots. The astrocyte feeder layer consisted of astrocytes cultured from postnatal day 0 (P0) pups. Cultures were maintained in Neurobasal medium supplemented with B27 (NB/B27) for 18-21 days before treatment with PrP^Sc^.

For omics-based analyses (transcriptomics, proteomics, and phosphoproteomics), which required larger number of neurons, the neuronal and astrocyte feeder layers were reversed. Hippocampal neurons were cultured at high density on poly-L-lysine-coated plates. After several hours, astrocytes plated on coverslips were inverted onto the neuron cultures. These cultures were maintained under the same medium conditions (NB/B27).

### Proteomic and phosphoproteomic analyses

Neurons were lysed in 8M Urea 50mM Triethylammonium bicarbonate (TEAB) buffer containing protease inhibitors (Sigma) and phosphatase inhibitors (Roche). After brief sonication on ice, the samples were reduced by addition of dithiothreitol (5 mM final concentration) for 60 min at room temperature and alkylated by the addition of iodoacetamide (5 mM final concentration) and incubation at room temperature for 30 min in the dark. Proteins were diluted with 50 mM TEAB to bring the urea concentration below 1 M, and digested with MS-grade trypsin (1:50 enzyme to protein ratio) at 37°C overnight followed by the addition of formic acid to a final concentration of 1%. The resulting peptides were desalted using a C18 Tips (Thermo Scientific) as per the manufacturer’s instructions.

Tryptic peptides were labeled with isobaric, tandem mass tags (TMT), followed by isolation of phosphopeptides on Fe-NTA beads (CubeBiotech). A portion (10%) of each sample prior to phosphopeptide isolation was saved for total proteome analysis. Briefly, peptides (90%) were resuspended in 800 µl of sample buffer consisting of 0.5% TFA and 80% ACN, and shaken at 1,000 rpm on a temperature-controlled shaker at 20°C for 15 min. Beads (10 µg per 100 µg of peptides) were washed with washing buffer consisting of 0.1% TFA and 80% ACN, then added to the resuspended peptides. Samples and beads were co-incubated for 30 min (shaken at 1000 rpm at 20°C). Bound phosphopeptides were washed twice with 800 µl of washing buffer and then serially eluted twice with 200 µl of 7.5% ammonium hydroxide and 50% ACN. The two eluates were combined and dried in a SpeedVac prior to LC/MS.

### Quantitative LC-MS analysis of peptides

LC-MS analysis was performed using a Q-Exactive HFX mass spectrometer(Thermo Scientific), interfaced with and EASY nLC 1200 ultra-high pressure pump system. Peptides were loaded onto a C18 pre-column (75 μm i.d. × 2 cm, 3 μm, 100 Å, ThermoFisher Scientific) and then separated on a capillary column (75 μm i.d. × 50 cm, 2 μm,100 Å, ThermoFisher Scientific) using gradient elution. The nLC flow rate was 250 nl/min. Peptides were injected and separated over a 90 min gradient. The mobile phase A was 0.1% FA-2% ACN-Water, and mobile phase B was 0.1% FA-80% ACN-Water. The gradient consisted of 2%-6% of mobile phase B for 5 min, was increased to 20% mobile phase B over 39 min, then increased to 35% over 23 min, and reached 98% over 3 min and maintained at 98% for 3 min before decreasing to 5% over 3 min. The MS instrument was operated in positive ion mode over a full mass scan range of m/z 350−1500 at a resolution of 120,000 with a normalized AGC target of 1e^5^. The source ion transfer tube temperature was set at 275°C and a spray voltage was set to 2.5kv. Data was acquired on data-dependent acquisition (DDA) mode. MS2 scans were performed at 45,000 resolution with a normalized collision energy of 33. The maximum injection time was set to 400ms. Dynamic exclusion was enabled using a time window of 45 s.

### Phosphoproteomic data analysis

MS2 spectra were processed and searched by MaxQuant (version 1.6.7) against a database containing native (forward) mouse protein sequences (UniProt) and reversed (decoy) sequences for protein identification. The search allowed for two missed trypsin cleavage sites, and phosphorylation (Ser/Thr/Tyr) was included as variable modification. Both carbamidomethylation of cysteine residues and TMT (peptide N-termini, K residues) were set as fixed modifications. Precursor ions were searched with a maximum mass tolerance of 4.5 ppm and fragment ions with a maximum mass tolerance of 20 ppm. The candidate peptide identifications were filtered, assuming a 1% FDR threshold based on searching the reverse sequence database. Quantification was performed using the TMT reporter on MS2. Proteins with less than 50% missing value across all samples were kept for quantification. The reporter ion intensities were log-transferred normalized. Bioinformatic analysis was performed in the R statistical computing environment (version 3.6) using Omics Notebook [73]. The p-value was obtained using moderated Student t-test (Limma) and multiple testing was controlled using Benjamini-Hochberg FDR method. Pathway enrichment was performed using GSEA [20] mapped to GO terms for Molecular Function, Biological Process, and Cellular Component Ontology.

### Phosphoproteomic data availability

Phosphoproteomic raw data are deposited in MassIVE under accession number MSV000100370.

### RNA-seq analysis

Total RNA was isolated from cultured neurons using the RNeasy Plus Mini Kit (Qiagen) following the manufacturer’s protocol. Briefly, samples were lysed in the provided RLT buffer containing 1% β-mercaptoethanol and homogenized using a sonicator. The lysate was then passed through an RNA-cleaning column to remove contaminants, and the RNA was eluted in

RNase-free water. The integrity and concentration of the isolated RNA were assessed using the High Sensitivity RNA Pico 6000 assay run on an Agilent 2100 Bioanalyzer (Agilent Technologies, USA). Only samples with RNA Integrity Numbers (RIN) greater than 8 were used for sequencing. RNA samples were then subjected to DNase I treatment (Thermo Fisher Scientific) to remove any contaminating genomic DNA, followed by another round of purification using the RNeasy column. RNA-seq was performed in-house at the Boston University Microarray & Sequencing Resource Core. Libraries were prepared using Illumina’s TruSeq Stranded mRNA Library Preparation kit according to the manufacturer’s protocol (Illumina, USA). To enrich for mRNA, 200 ng of total RNA for each sample was incubated with magnetic Oligo-(dT) beads. Enriched mRNA was fragmented at 94°C for 8 minutes. First strand cDNA was generated using random primers, and incorporation of dUTP occurred during synthesis of second strand cDNA to indicate strandedness. Double-stranded cDNA then underwent A-tailing and adaptor ligation. Sample-specific multiplex indices were incorporated during PCR amplification (15 cycles) according to the manufacturer’s protocol (Illumina, USA). Size distribution and molarity of amplified cDNA libraries were assessed via the Bioanalyzer High Sensitivity DNA Assay (Agilent Technologies, USA). All cDNA libraries were sequenced on an Illumina NextSeq 500 instrument (Illumina, USA) targeting 30-40 million reads per sample (75 x 75 paired-end reads). The resulting sequencing data were processed and analyzed using standard bioinformatic pipelines. Pathway enrichment was performed using GSEA [20] mapped to GO terms for Molecular Function, Biological Process, and Cellular Component Ontology.

### Chemo-omic analysis using L1000 and P100 databases

Transcriptomic and proteomic data were utilized as inputs for a systematic multi-platform connectivity analysis pipeline to identify candidate therapeutic compounds modulating PrP^Sc^-induced perturbations. RNA-seq raw counts were processed to generate normalized expression values, followed by differential gene expression analysis at each time point (0.5, 2, 4, and 24 hours). Unfiltered DGE values, separated into up- and down-regulated genes (as listed in Table S4), were used as inputs into the L1000 database. Proteomic and phosphoproteomic datasets were analyzed to determine proteins exhibiting significant abundance or phosphorylation changes (p < 0.05), which were compiled as filtered input feature lists. These gene and protein feature sets were harmonized to recognized gene symbols and annotations compatible with downstream connectivity tools. Values for differentially phosphorylated proteins (p<0.05, separated into up- and down changes; see Table S2) were used as inputs into the P100 database.

For transcriptomic perturbational connectivity, two independent platforms were employed to comprehensively capture candidate compound signatures: (1) the CLUE platform (Connectivity Map Linked User Environment; https://clue.io/) accessed through Metascape [74], which calculates similarity metrics based on both the magnitude and direction of expression changes across the landmark L1000 gene set, thus capturing nuanced expression pattern alignments; and (2) L1000CDS² [75], which uses directional gene set enrichment to identify compounds predicted to reverse the up/down regulation pattern independent of fold change magnitude, emphasizing functional reversal potential. Both tools query the Library of Integrated Network-Based Cellular Signatures (LINCS) L1000 dataset [16–19] comprising thousands of small-molecule perturbations.

In parallel, the phosphoproteomic signatures were mapped to the LINCS P100 phosphoproteomic database using CLUE, enabling compound connectivity inference based on phosphorylation state modulation rather than transcriptomic profiles. This complementary analysis aimed to identify compounds capable of reversing phosphorylation changes induced by PrP^Sc^.

Connectivity results from all platforms were integrated by identifying compounds recurrently predicted with high confidence (top connectivity scores or enrichment metrics) across multiple time points, genotypes, and data modalities. Candidates shared by both transcriptomic and phosphoproteomic analyses were prioritized for enhanced confidence.

Finally, the resulting shortlist of compounds underwent manual curation to assess biological plausibility in the context of neurodegeneration and prion pathology, confirmation by prior literature, and practical considerations such as commercial availability and suitability for experimental follow-up. This rigorous pipeline ensured that candidate therapeutic agents stemmed from robust, statistically grounded molecular signatures and cross-validated connectivity predictions, providing a rational basis for downstream testing.

### Immunocytochemistry and microscopic imaging

Cells grown on glass coverslips were washed with phosphate-buffered saline (PBS) and fixed in 4% paraformaldehyde for 12 minutes at room temperature. For detection of intracellular antigens, cells were permeabilized with 0.1% Triton X-100 in PBS for 5 minutes prior to blocking. Non-specific binding was blocked with 1% bovine serum albumin (BSA) in PBS for 30 minutes at room temperature.

Neurons were stained with the primary antibodies listed in Table S6 and with fluorescent phalloidin (labeled with Alexa Fluor 488 or rhodamine) to label F-actin in dendritic spines. Antibodies and phalloidin were diluted in 1% BSA/PBS and applied to cells for 2 hours at room temperature. After washing with PBS, fluorophore-conjugated secondary antibodies diluted in 1% BSA/PBS were applied for 1 hour at room temperature in the dark. Cells were then washed five times (5 minutes each) in PBS and counterstained with DAPI (Invitrogen) to visualize nuclei. Coverslips were mounted using Vectashield Mounting Medium (Vector Laboratories) and stored at 4°C until imaging.

Fluorescence imaging was performed using Zeiss laser-scanning confocal microscopes with identical acquisition settings across conditions. For experiments to quantitate dendritic spines, images were acquired using a Zeiss LSM 700 microscope. For localization of proteins within dendritic spines and neuronal somata, images were acquired using a Zeiss LSM 880 microscope. A 63x oil immersion objective (numerical aperture, NA=1.4) was used for high-resolution image capture.

### Dendritic spine quantitation

Dendritic spine density was assessed to evaluate synaptotoxic effects, as previously described [14, 15]. For quantification, 4-5 isolated dendritic segments were selected per image. Images were processed in ImageJ (FIJI) using a threshold optimized to include dendritic spine outlines while excluding background fluorescence [76]. Spine numbers were normalized to the corresponding dendritic length and reported as spines per micrometer (spines/μm). For each condition, 15-24 neurons were analyzed across 3-4 independent experiments.

### Quantitation of immunofluorescence intensity

Fluorescence staining intensity of neuronal proteins (CaMKII, PKC, GSK3β, PSD95, NR2B) within defined regions of interest (ROIs) was quantified using ImageJ (FIJI). For each sample, ROIs were manually selected based on the area of interest, such as dendritic spines, dendrites, neuronal somata, or intracellular compartments. Fluorescence measurements were obtained by setting the analysis parameters to include Area and Integrated Density to calculate the total signal within the selected ROI.

Each image was processed to ensure uniformity, including background subtraction and optimization of thresholding to exclude non-specific fluorescence signals. The mean fluorescence intensity per pixel was calculated by dividing the integrated density by the ROI area. To control for variability between experimental conditions, all fluorescence intensities were normalized to the average intensity of the control group. Data from at least 10 randomly selected fields per condition were quantified to ensure robust sampling. For each neuron, 2-3 isolated ROIs were selected. For each condition, 15-24 neurons from 3-4 independent biological replicates were analyzed.

### Western blotting

Neuronal lysates were prepared in RIPA buffer. Total protein concentration was quantified using a bicinchoninic acid (BCA) assay (Pierce). Equal amounts of protein (typically 20-30 μg) were mixed with an appropriate volume of SDS-PAGE loading buffer and heated for 5 minutes at 99°C prior to separation by SDS-PAGE using Tris-Glycine gels. Following electrophoresis, proteins were transferred onto Immobilon PVDF membranes (Millipore, Immobilon-P). Membranes were blocked for 30 minutes with 5% BSA in TBS buffer and subsequently incubated overnight at 4°C with primary antibodies. Antigen-antibody complexes were detected using HRP-conjugated secondary antibodies followed by development with ECL (Millipore) and visualization with a Bio-Rad Chemidoc XRS Molecular Imager; or using fluorescent secondary antibodies from LI-COR Biosciences (IRDye 800 (green) or IRDye 680 (red) for goat anti-mouse; IRDye 680 (red) for goat anti-rabbit), followed by visualization using the LI-COR Odyssey Classic Infrared Imaging System.

### Statistical analysis

All quantification was performed in at least three independent biological replicates. Statistical comparisons were conducted using two-tailed, unpaired, Student’s t-tests (GraphPad Prism). Data are presented as bar graphs showing meanc±cstandard error of the mean (SEM), or as violin plots with the median indicated by a dotted red line. Details regarding the number of replicates (n), type of statistical test used, and significance levels are provided in the corresponding figure legends.

## Author Contributions

**N.T.T.L.** (Conceptualization, Formal Analysis, Funding Acquisition, Investigation, Methodology, Validation, Visualization, Writing-Original Draft, Review & Editing); **R.C.C.M.** (Conceptualization, Investigation, Methodology, Visualization, Writing-Review and Editing); **C.F.** (Investigation, Methodology); **A.S.** (Formal Analysis); **A.T.L.** (Formal Analysis); **W.L.** (Formal Analysis, Writing-Review and Editing); **J.K.** (Formal Analysis, Investigation); **B.B.** (Formal Analysis, Software, Visualization); **A.E.** (Conceptualization, Funding Acquisition); **D.A.H.** (Conceptualization, Formal Analysis, Funding Acquisition, Methodology, Supervision, Visualization, Writing-Original Draft, Review & Editing)

## Acknowledgements

This work was supported by NIH grants R01NS065244 and R21NS107755 (to D.A.H.); and R01AG064932 and RF1AG061706 to A.E. N.T.T.L. was supported by a Warren Alpert Distinguished Scholar Award. We thank the Microarray and Sequencing Resource Core at Boston University Chobanian & Avedisian School of Medicine for performing RNA-seq analysis, and Michael Blower for transcriptomic bioinformatic support.

## SUPPLEMENTARY TABLE LEGENDS

**Table S1. Differentially expressed proteins from proteomic analysis of neurons after 1 h of PrP^Sc^ treatment.**

**Table S2. Differentially phosphorylated proteins from proteomic analysis of neurons after 1 h of PrP^Sc^ treatment.**

**Table S3. Pathway enrichment analysis of phosphoproteomic data from neurons after 1 h of PrP^Sc^ treatment.**

**Table S4. Differential gene expression from RNA-seq analysis of neurons after 0.5, 2, 4, and 24 h of PrP^Sc^ treatment.**

**Table S5. Pathway enrichment analysis of RNA-seq data from neurons after 0.5. 2, 4, and 24 h of PrP^Sc^ treatment.** Pathways were filtered for adjusted p-values <0.05, and the top 20 pathways were then ranked based on NES.

**Table S6. Antibodies used in this study.**

## SUPPLEMENTARY FIGURE LEGENDS

**Figure S1. Time course of PrP^Sc^-induced dendritic spine retraction and experimental plan for collection of samples. (A)** Mature hippocampal neurons (21DIV) were treated with purified PrP^Sc^ for 30 min, 1 hour, 2 hours, 4 hours, or 24 hours, or with mock-purified material from uninfected brains for 24 hours. Neurons were then stained with fluorescent phalloidin to reveal dendritic spine morphology. Scale bars = 5 μm. **(B)** Quantification of spine number in PrP^Sc^-treated neurons compared to mock-treated cultures. Pooled measurements were collected from 5-7 neurons and 15-25 dendritic regions from 3 independent experiments. Data are shown as the mean ± SEM. Statistical analysis was performed using unpaired t-tests. Significance is indicated as: *p < 0.05, ***p < 0.001. **(C)** Experimental set-up for RNA-seq and phosphoproteomic analysis.

**Figure S2. Proteomic and phosphoproteomic analysis of hippocampal neurons treated with PrP^Sc^. (A)** Principal component analysis (PCA) of proteomic and phosphoproteomic data from five biological replicates of hippocampal neurons treated for 1 hr with PrP^Sc^ (PRP, cyan dots) or material that was mock-purified from uninfected brains (NBH, red dots). **(B)** Venn diagram comparing the number of proteins identified in the proteomic and phosphoproteomic analyses. The numbers refer to proteins that were identified based on at least two peptides. **(C)** Volcano plots depicting proteomic and phosphoproteomic data. Cut-offs are │log2fold- change│>0.25 and adjusted p-value <0.05. **(D)** Heat maps of proteomic and phosphoproteomic changes with Z-scores in the top 1% mapped onto corresponding genes.

**Figure S3: Volcano plots from transcriptomic analysis of hippocampal neurons treated with PrP^Sc^.** Neurons were treated with purified PrP^Sc^ for 30 min **(A)**, 2 hr **(B)**, 4 hr **(C)**, and 24 hr **(D)**. Cut-off values were |log2fold change│>0.25 and p-value <0.05.

**Figure S4. Screening of inhibitors from the chemogenomics pipeline for their ability to prevent spine retraction.** Hippocampal neurons were pretreated with non-toxic compounds listed in Table 1 at 250 nM, then exposed to PrP^Sc^ or mock-purified material for 24 hrs. After fixation, neurons were stained with Alexa 488-labeled phalloidin for visualization of dendritic spines. The top pair of panels shows neurons that were exposed to PrP^Sc^ or mock-purified material in the absence of any compound. White and red arrowheads in the top panels indicate healthy and retracted dendritic spines, respectively. Scale bars = 5 μm. Compounds and targets are color-coded as indicated in the legend and in Table 1.

**Fig S5: Western blot analysis of total and phosphorylated forms of CaMKII, PKC, and GSK3**β **in hippocampal neurons treated with PrP^Sc^.** Western blots of neuronal lysates were probed with the indicated antibodies to total and phosphorylated forms of CaMKII **(A)**, PKC **(D)**, and GSK3β **(F)**. Actin was used as a loading control in panels D and F. Blots were quantified by Image J **(B, C, E, G)**. Data are shown as the mean ± SEM. Statistical analysis was performed using unpaired t-tests. Significance is indicated as: ns (not significant).

**Figure S6: CaMKII inhibitor KN93 prevents translocation of CaMKII to dendritic spines. (A)** Hippocampal neurons were pre-treated with KN-93 for 2 hours, then treated for 1 hour with either mock-purified material (Mock) or purified PrP^Sc^ (PrP^Sc^). Cultures were then fixed and stained with fluorescent phalloidin (gray), along with antibodies to total CaMKII (red) and N-methyl D-aspartate receptor subtype 2B, NR2B (green). Dotted lines in the smaller panels outline the positions of intact spines, based on phalloidin staining. Scale bars = 5 μm. Violin plots show quantitation of Total CaMKII **(B)**, Total NR2B **(C)**, and CaMKII/NR2B **(D)** in spine regions. Measurements were collected from 5-7 neurons, and 15 dendritic regions from at least 2 independent experiments. Each data point for the PrP^Sc^-treated samples was normalized to the average Mock value from the same experiment. Dotted red lines in each violin plot indicate the median. Statistical analysis was performed on SEM values using unpaired t-tests. Significance is indicated as: ns (not significant), *p < 0.05, **p < 0.01, ****p < 0.0001.

**Figure S7: PKC inhibitor Ro 32-0432 prevents accumulation of primed PKC in endosomes within the soma and in dendritic spines.** Hippocampal neurons were pre-treated with Ro 32-0432 hydrochloride for 2 hours and were then treated either with mock-purified material **(A)**, or with purified PrP^Sc^ for 30 min **(B)**, 2 hours **(C)**, 4 hours **(D)** or 24 hours **(E)**. After fixation, neurons were stained with fluorescent phalloidin (gray), along with antibodies to total PKC (green) and PKC-pS660 (red). Boxed regions in the square panels are shown at higher magnification in the rectangular panels below. Dotted lines in the higher magnification panels outline the positions of intact spines, based on phalloidin staining. Asterisks mark the locations of neuronal somata. Scale bars = 5 μm. Violin plots show quantitation of pS660/total PKC ratios within somata **(F)** and spines **(G)**. Measurements were collected from 5-7 neurons, 30-45 dendritic and somatic regions from at least 2 independent experiments. Each data point for the PrP^Sc^-treated samples was normalized to the average Mock value from the same experiment. Dotted red lines in each violin plot indicate the median. Statistical analysis was performed on SEM values using unpaired t-tests. Significance is indicated as: ns (not significant), **p < 0.01.

