## Supplemental Figures for "Chemo-omic pipeline enables discovery of prion synaptotoxic pathways and inhibitory drugs"

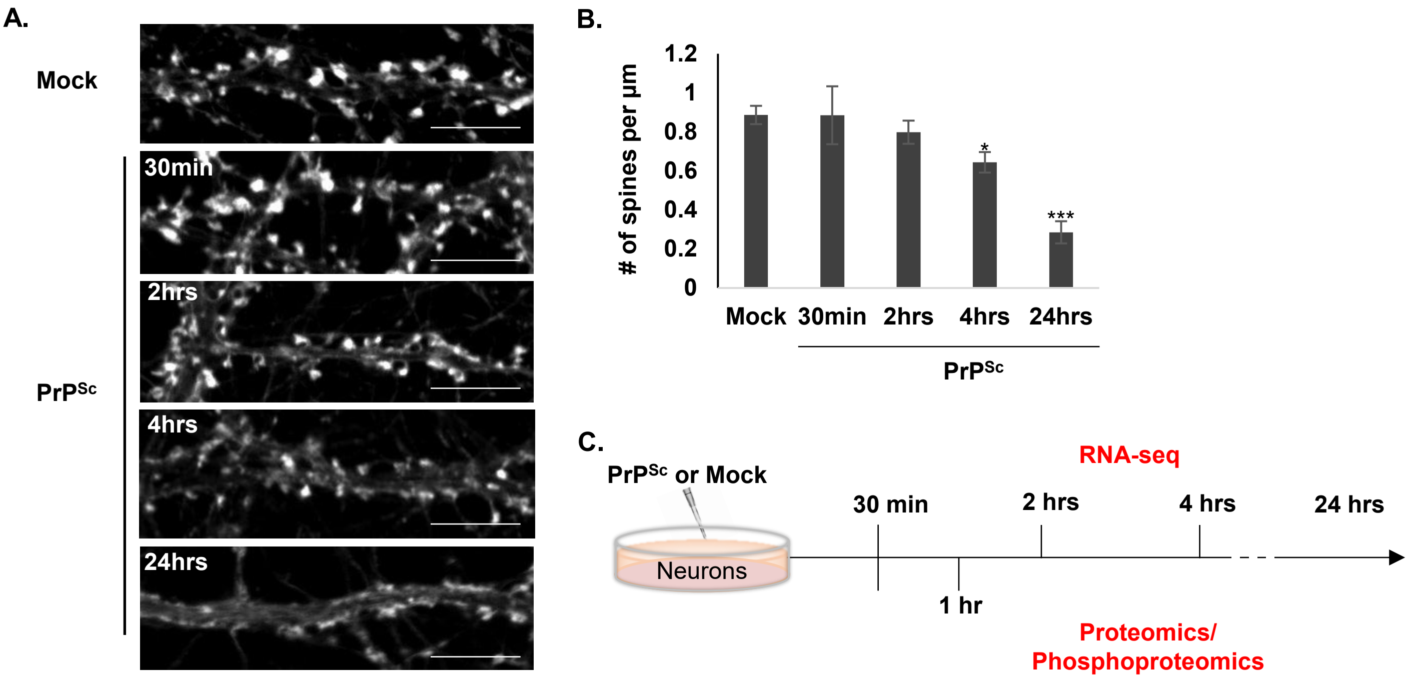


**FIGURE S1 : Time course of PrP$^{Sc}$-induced dendritic spine retraction and experimental plan for collection of samples**

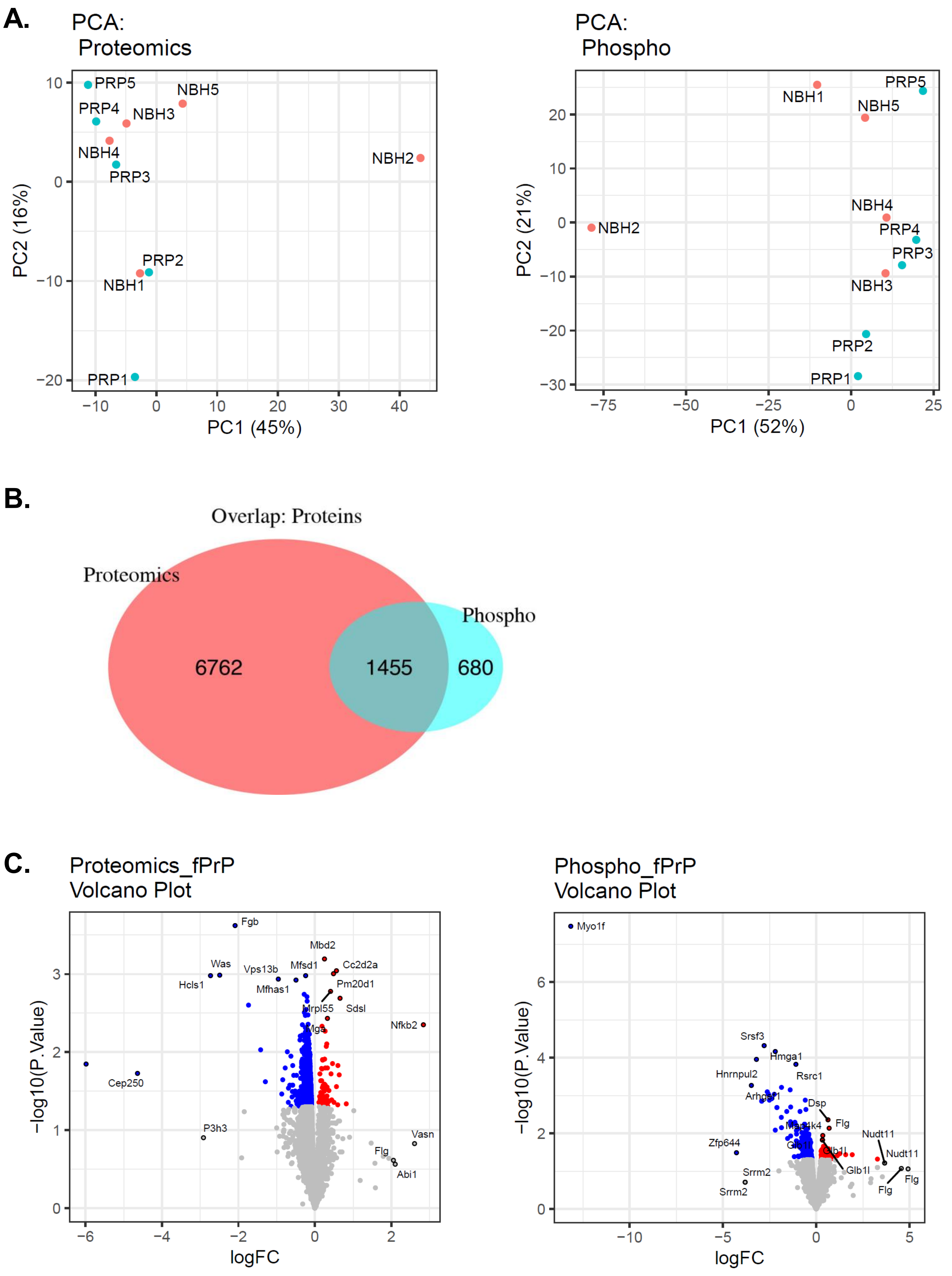


**FIGURE S2: Proteomic and phosphoproteomic analysis of hippocampal neurons treated with PrP$^{Sc}$**

D.

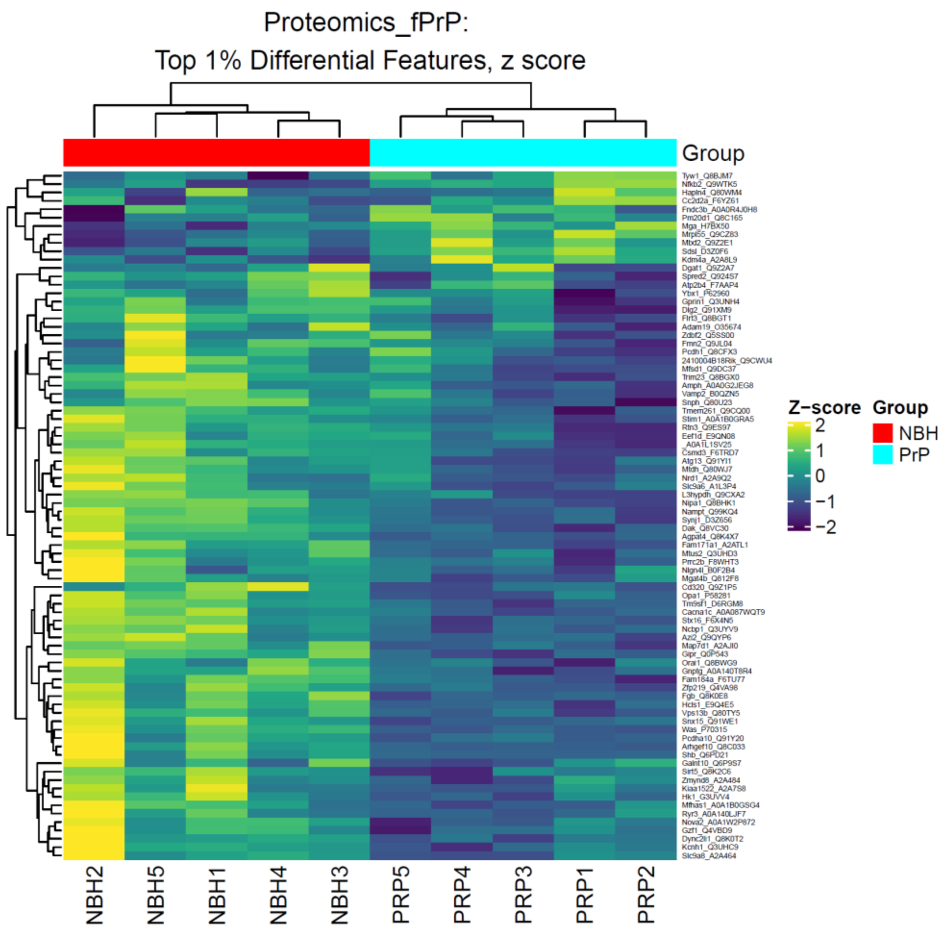


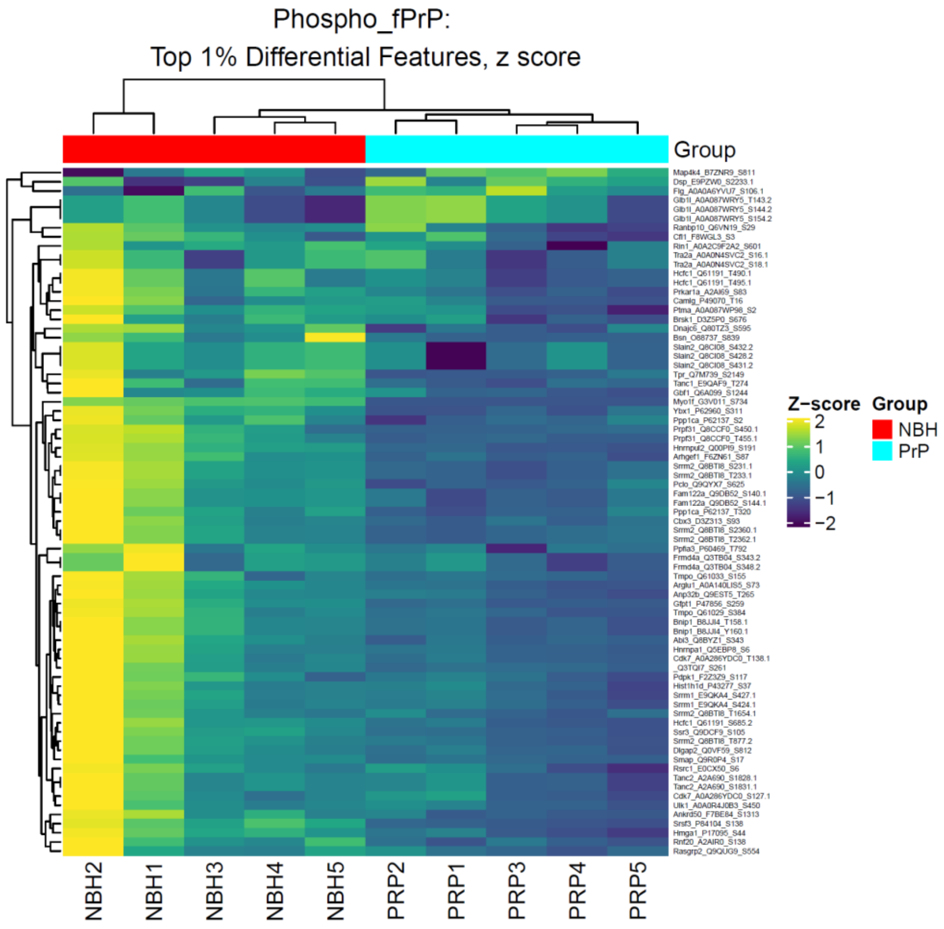


**FIGURE S2 (con’t): Proteomic and phosphoproteomic analysis of hippocampal neurons treated with PrP$^{Sc}$ for 1 hour**

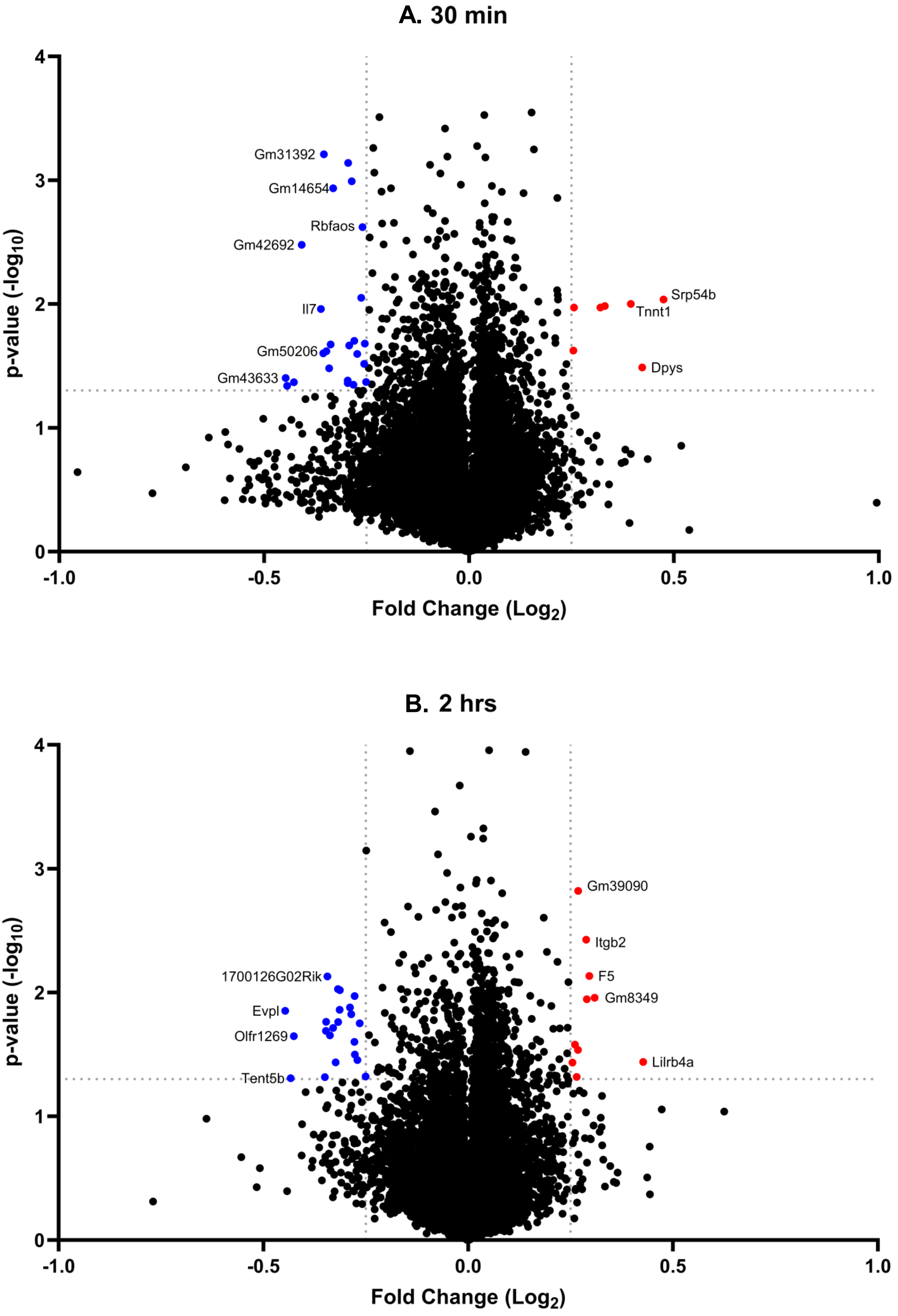


**FIGURE S3: Volcano plots from transcriptomic analysis of hippocampal neurons treated with $PrP^{Sc}$**

#### C. 4 hrs

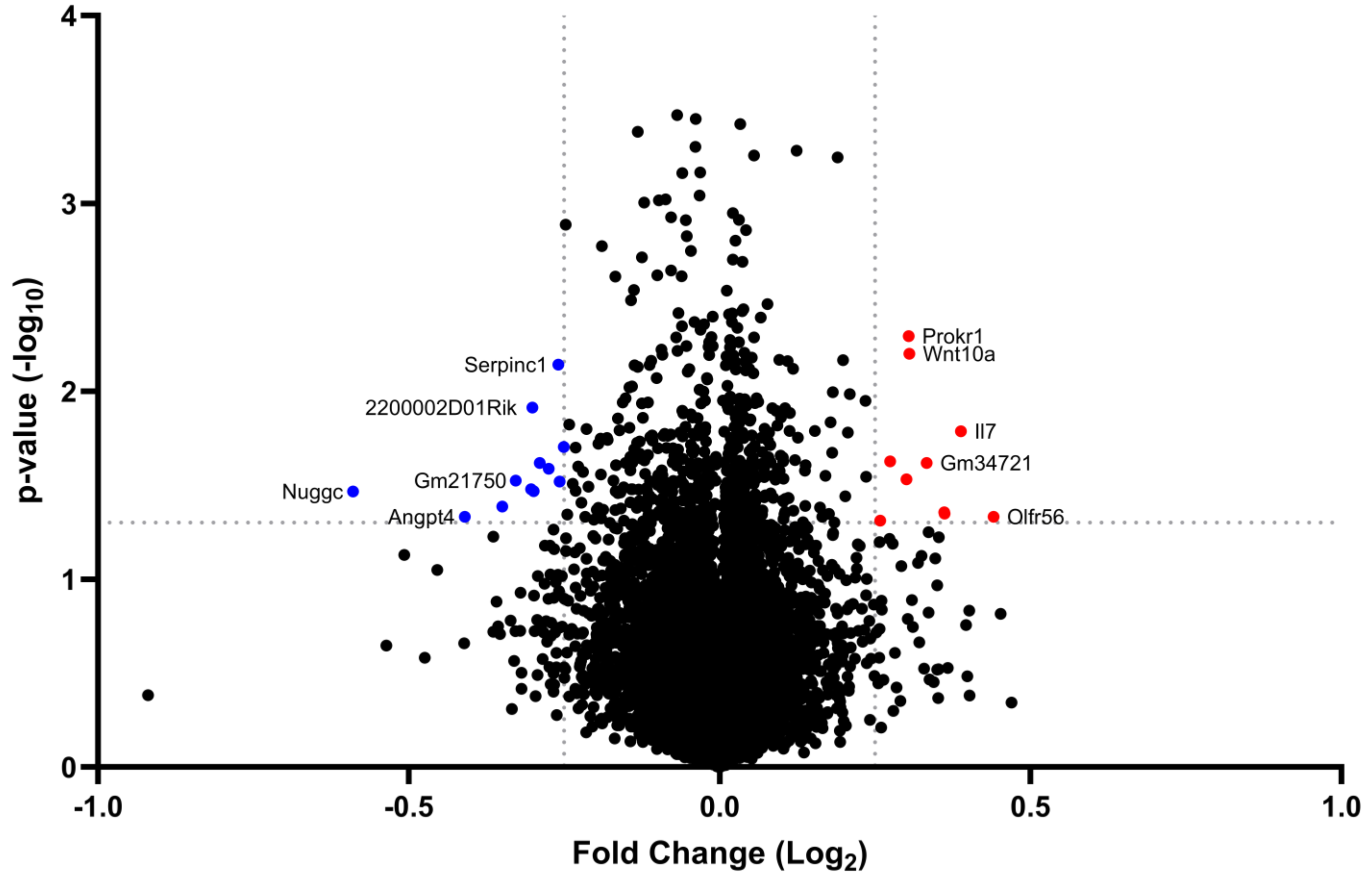


#### D. 24 hrs

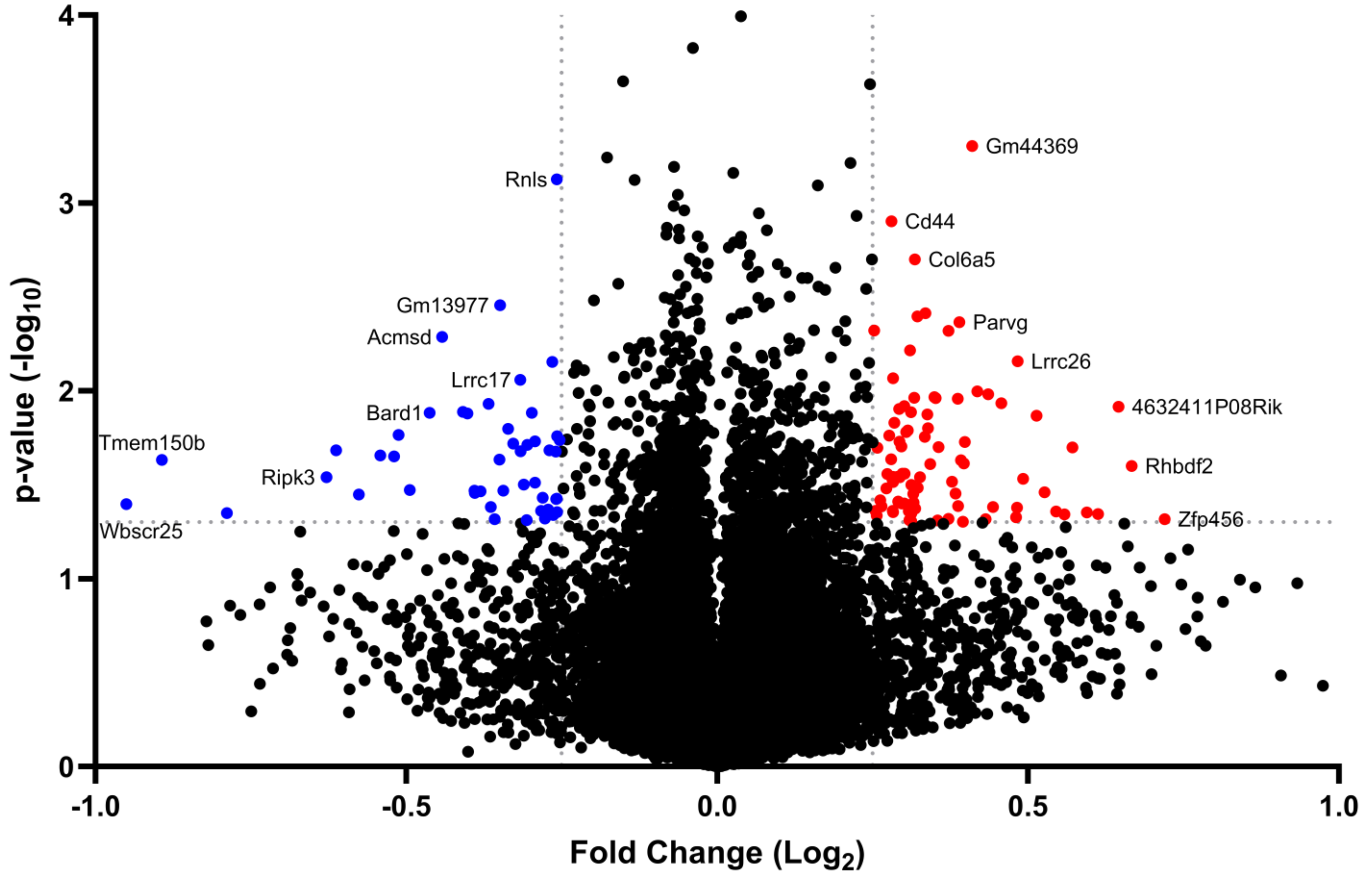


**FIGURE S3 (con't): Volcano plots from transcriptomic analysis of hippocampal neurons treated with $PrP^{Sc}$**

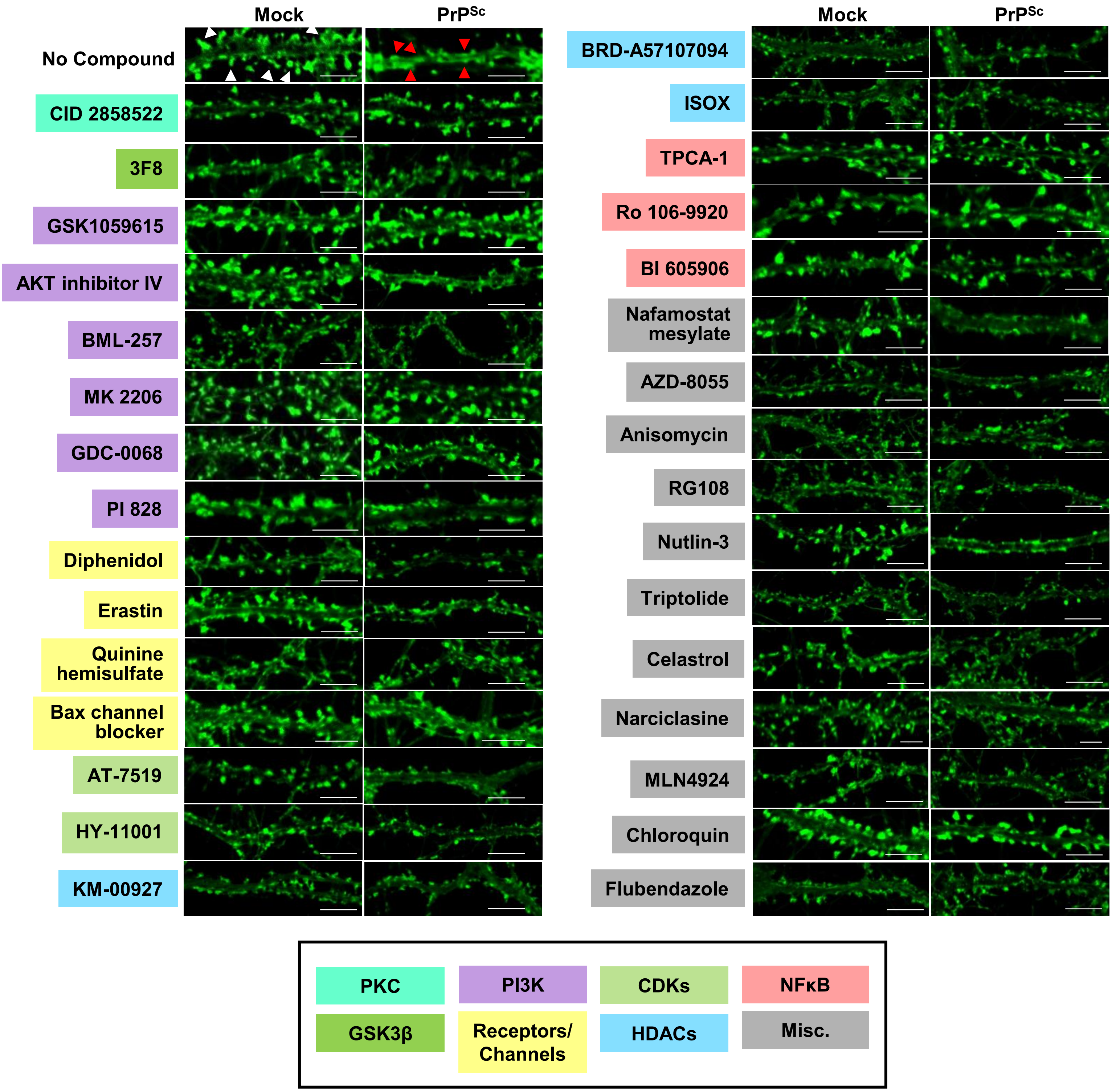


**FIGURE S4: Screening of inhibitors from the chemogenomics pipeline for their ability to prevent spine retraction**

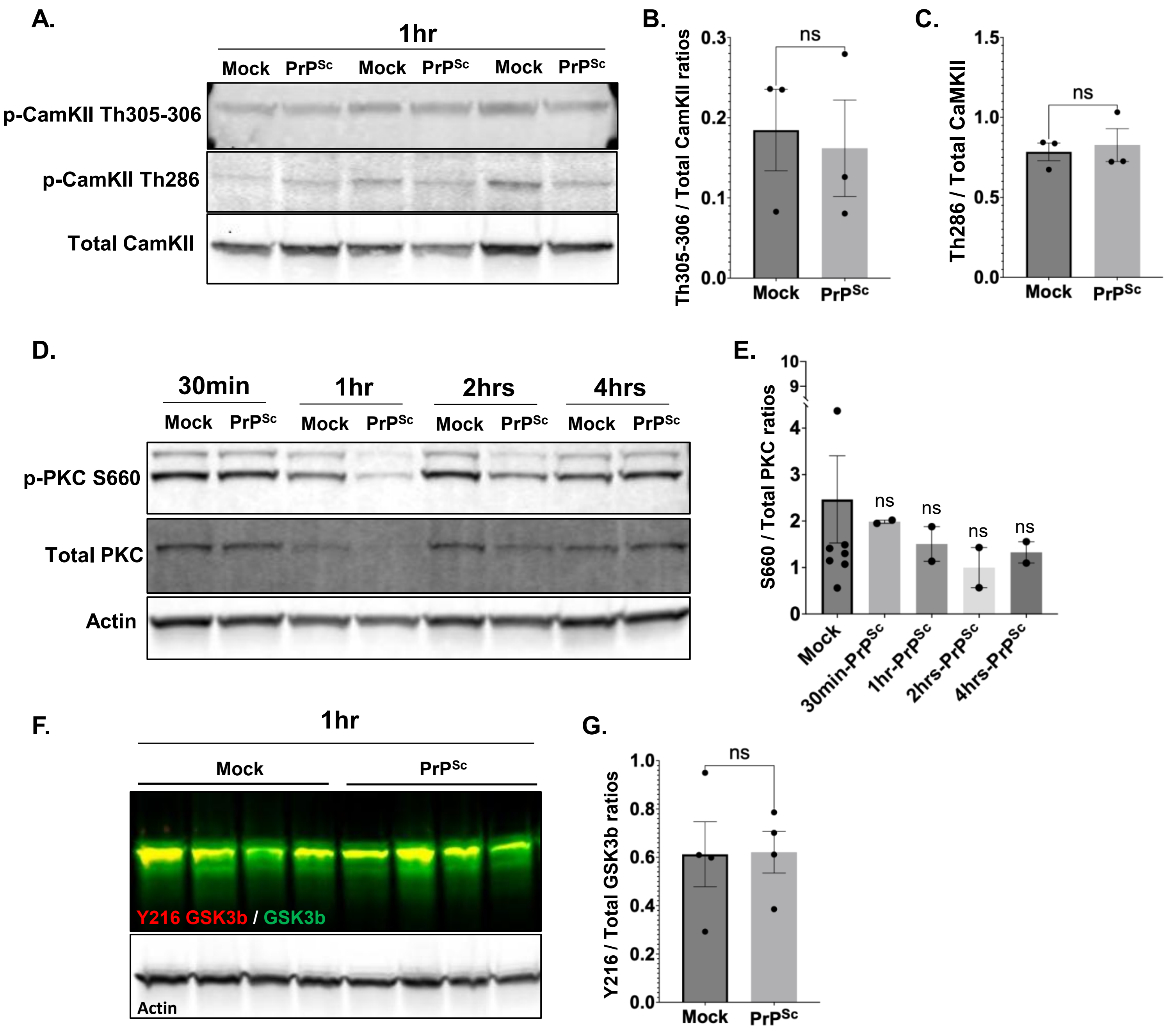


### Figure S5: Western blot analysis of total and phosphorylated forms of CamKII, PKC, and GSK3β in hippocampal neurons treated with $PrP^{Sc}$

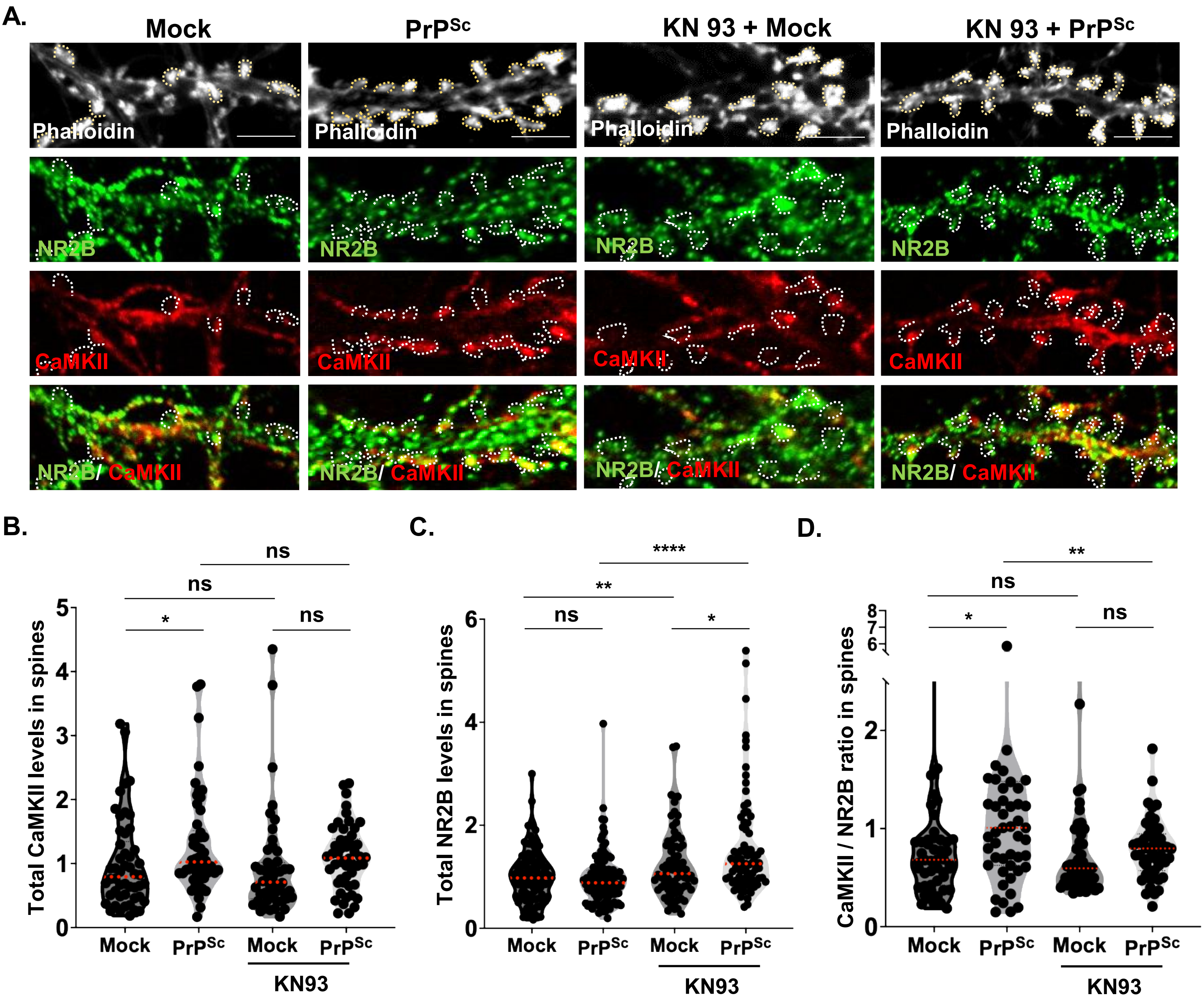


### FIGURE S6: CaMKII inhibitor KN93 prevents translocation of CaMKII to dendritic spines.

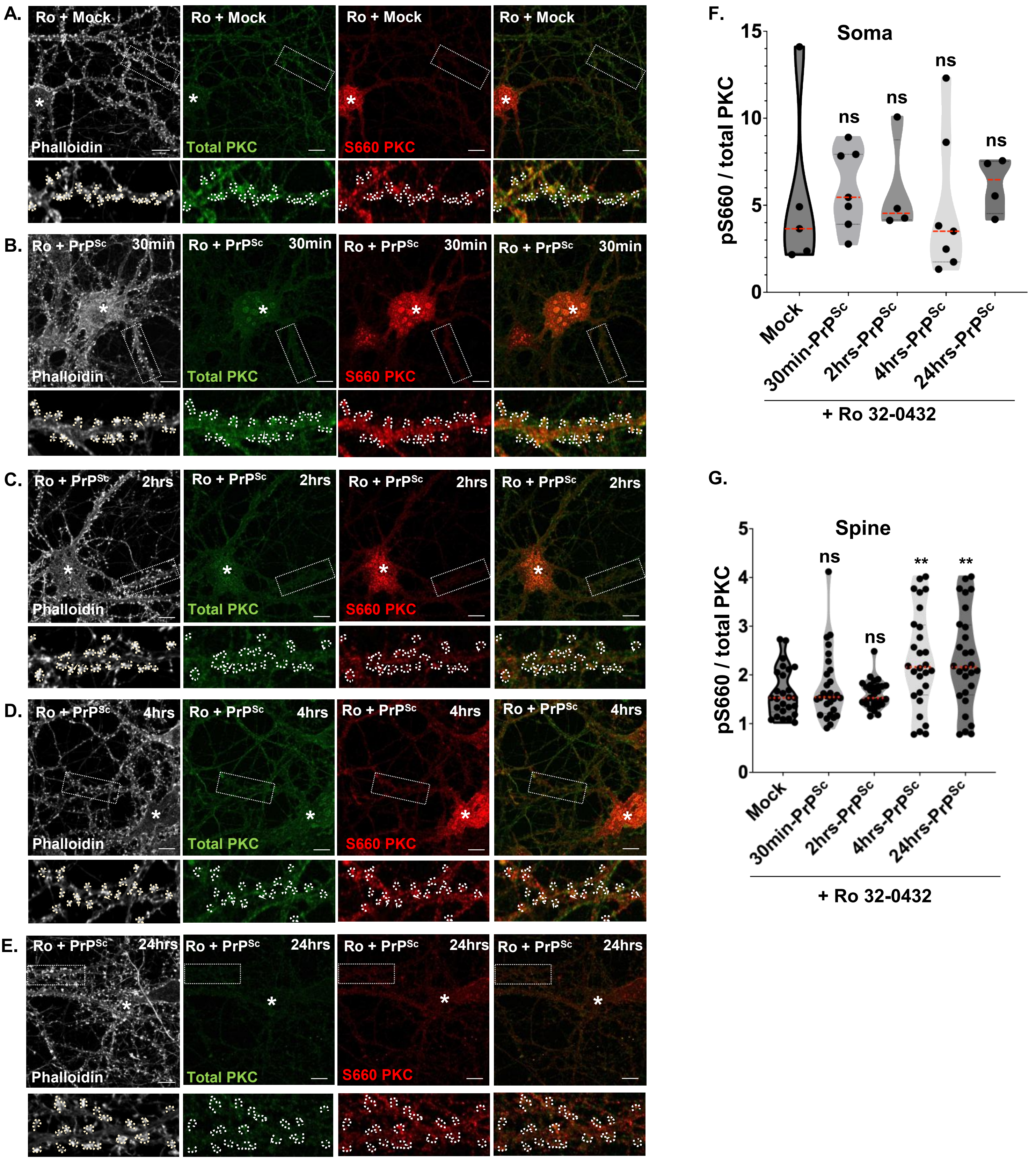


**FIGURE S7: PKC inhibitor Ro 32-0432 prevents accumulation of primed PKC in endosomes within the soma and in dendritic spines.**
